# A Trisomy 21 Model Atlas reveals conserved dosage effects and context-specific transcriptional responses across mouse and human models of Down syndrome

**DOI:** 10.64898/2026.09.20.751055

**Authors:** Karen Rossmassler, Ariel Timkovich, Brian Niemeyer, Kyndal Schade, Lauren Dunn, Macallister C. Harris, R. Adam Harris, Srija Chillamcherla, Angela L. Rachubinski, Joaquin M. Espinosa, Kelly D. Sullivan, Matthew D. Galbraith

## Abstract

Trisomy 21, the genetic basis of Down syndrome, causes widespread transcriptional dysregulation with tissue-specific phenotypic consequences, but how this propagates across tissues, developmental stages, and model systems remains incompletely understood. We present the initial release of the Trisomy 21 Model Atlas, comprising bulk transcriptomic profiling and histopathology across seven tissues and three developmental timepoints in the Dp(16)1Yey mouse model along with transcriptional profiles from human iPSC-derived cell types. Triplicated genes show consistent, largely dosage-proportional increases in expression across tissues, developmental stages, sexes, and species. Nontriplicated genes, which account for the substantial majority of differential expression, are instead highly context-specific. Inflammatory and interferon-related gene sets are broadly enriched yet composed of distinct genes in each context, and integration with histopathology links expression signatures to tissue-specific pathological features. The Atlas is openly accessible and designed for expansion, providing a framework for interpreting conserved and context-specific consequences of chromosome 21 gene dosage.

## INTRODUCTION

Down Syndrome (DS) is a complex, multi-organ condition driven by dysregulated gene expression caused by trisomy of chromosome 21 (T21). T21 affects the development and/or function of nearly every organ system, with key impacted tissues including the brain, heart, lung, liver, intestine, immune system, bone marrow, kidney, skin, and muscle^1^. Consequently, individuals with DS are predisposed to a broad range of co-occurring conditions that arise both directly and through their cumulative effects during development and across the lifespan^1–5^, including neurological (cognitive impairment, autism spectrum disorder, Alzheimer’s disease)^5,6^, cardiopulmonary (congenital heart defects, airway malformations, recurrent respiratory infections)^1,7–9^, and immune and hematologic (autoimmune disorders, leukemias)^10,11^, among others^12–18^.

Although altered chromosome 21 gene dosage clearly contributes to dysregulation of gene expression and downstream signaling, how these effects manifest across tissues, and the specific mechanisms linking T21 to the many developmental and clinical hallmarks of DS, are not yet fully understood. Comprehensively mapping the impacts of T21 on gene expression and organ development is therefore critical, both to identify individual genes acting as contributors or modifiers of specific phenotypes – and thus potential therapeutic targets – and to build a more complete picture of DS pathophysiology as a whole^19^.

Much of our current understanding of T21-driven pathophysiology in humans is drawn from a small number of readily accessible sample types, chiefly peripheral blood, leaving the downstream consequences of T21 in many organs and developing tissues incompletely characterized. Directly sampling tissues in humans is challenging: many organs cannot be safely or ethically biopsied from individuals who are otherwise relatively healthy, and collection of post-mortem or surgical specimens is laborious, sporadic, and typically limited to a single time point per individual. Critically, even a complete set of human tissue samples would remain largely observational, lacking the controlled experimental manipulations required for establishing cause-and-effect relationships between chromosome 21 gene dosage and downstream phenotypes. Model systems therefore remain indispensable: they extend molecular characterization to tissues and developmental windows that are inaccessible in humans, and they enable the experimental interventions required to address mechanistic hypotheses.

In mice, genes orthologous to those encoded on HSA21 are distributed across MMU10, 16, and 17, rather than a single chromosome. This genomic architecture has led to a proliferation of mouse models, each capturing varying degrees of orthology and synteny with HSA21, and in some cases additional non-orthologous genes, rather than a single complete recapitulation of the human trisomy^20^. These include Ts65Dn, Ts66Yah, Dp(16)1Yey, and TcMAC21, among others, each with its own trade-offs in terms of coverage, genetic background, and breeding characteristics^21–23^. The Dp(16)1Yey (henceforth Dp16) model used in this study carries a segmental duplication of the MMU16 region syntenic to the largest cluster of HSA21 genes and recapitulates multiple key hallmarks of DS, including abnormal craniofacial development, congenital heart defects, cognitive impairment, altered hematopoiesis, and immune dysregulation^22,24,25^.

Together, mouse models and human iPSC-derived systems provide complementary tools to interrogate the impacts of T21 on gene expression, each with distinct advantages and limitations: mice offer intact organismal physiology, tissue architecture, and developmental staging that remain difficult to replicate *in vitro*, while iPSC-derived systems allow direct examination of dosage effects in a human genetic background and access to otherwise hard-to-obtain human tissue types. Realizing the full potential of these tools requires not only characterizing transcriptomic and pathophysiological changes across tissues, cell types, and developmental stages, but also enabling systematic comparison of findings across model systems. Multi-tissue transcriptomic data from Dp16 mice have been reported in the context of specific phenotypes^26^, but a resource designed for cross-tissue, cross-stage, and cross-species comparison has not previously been available. The Trisomy 21 Model Atlas is designed to build exactly this: a comprehensive resource, readily accessible to the research community, spanning tissues, developmental stages, and mouse and human model systems side by side. The Atlas combines bulk transcriptomic profiling with histological analysis, linking tissue- and cell-specific gene expression changes directly to organ pathology and architecture. It further incorporates five differentiated cell types derived from a shared panel of trisomic and disomic human iPSCs, spanning all three germ layers and profiled by RNA sequencing. This extends prior work that profiled iPSCs and derived neural progenitor cells by microarray^27^, and enables direct comparison of *in vitro* human systems to mouse models of DS.

Here, we present the initial data release of the Trisomy 21 Model Atlas, spanning seven tissues and three developmental timepoints in the Dp16 mouse model together with transcriptional profiles from multiple human iPSC-derived cell types. Across this dataset, triplicated genes show remarkably consistent, dosage-proportional upregulation in essentially every tissue, timepoint, and cell type examined: a conserved molecular signature of T21 that holds across mouse and human systems. In sharp contrast, the much larger set of nontriplicated genes that respond to trisomy is overwhelmingly tissue-and cell-type-specific, indicating that the downstream consequences of chromosome 21 gene dosage are shaped as much by biological context as by the trisomy itself. Sex-stratified analyses suggest a similar pattern: dosage-dependent upregulation of triplicated genes is largely preserved between males and females, while nontriplicated transcriptional responses show greater sex-specific variability, adding a further axis of biological context to the Atlas. At the pathway level, inflammatory and interferon signaling emerge as a shared theme across nearly all contexts, yet are composed of distinct sets of genes in each context. Common phenotypic themes in DS can therefore arise from divergent underlying molecular responses. Together, these findings establish a foundational, cross-model view of T21-driven gene dysregulation.

The Atlas makes these data openly available: all datasets described herein are freely available as a dedicated collection through the Experimental Models of Down Syndrome portal (experimentalmodels.includedcc.org) within the INCLUDE Data Hub, where researchers can search by organ, cell type, or gene to examine how T21 affects their pathway or process of interest across multiple mouse and human model systems. This initial release represents a foundation rather than an endpoint: additional mouse models, cell lines, and data modalities, including single-cell transcriptomics and multiplexed imaging, are already in progress and will be incorporated in future Atlas releases. We anticipate that this growing, openly accessible resource will be of value to established and new investigators alike, thus lowering the barrier to entry for the DS research community while providing common ground for cross-model comparison and hypothesis generation.

## RESULTS

### Overview of the Trisomy 21 Model Atlas

The Trisomy 21 Model Atlas currently comprises bulk transcriptomic profiling and histopathology across seven tissues and three developmental timepoints in Dp16 and WT mice, together with transcriptional profiles from undifferentiated iPSCs and five differentiated human iPSC-derived cell types, all from a shared panel of trisomic and disomic donors (**Figure 1A, Table S1**).

**Figure 1.**
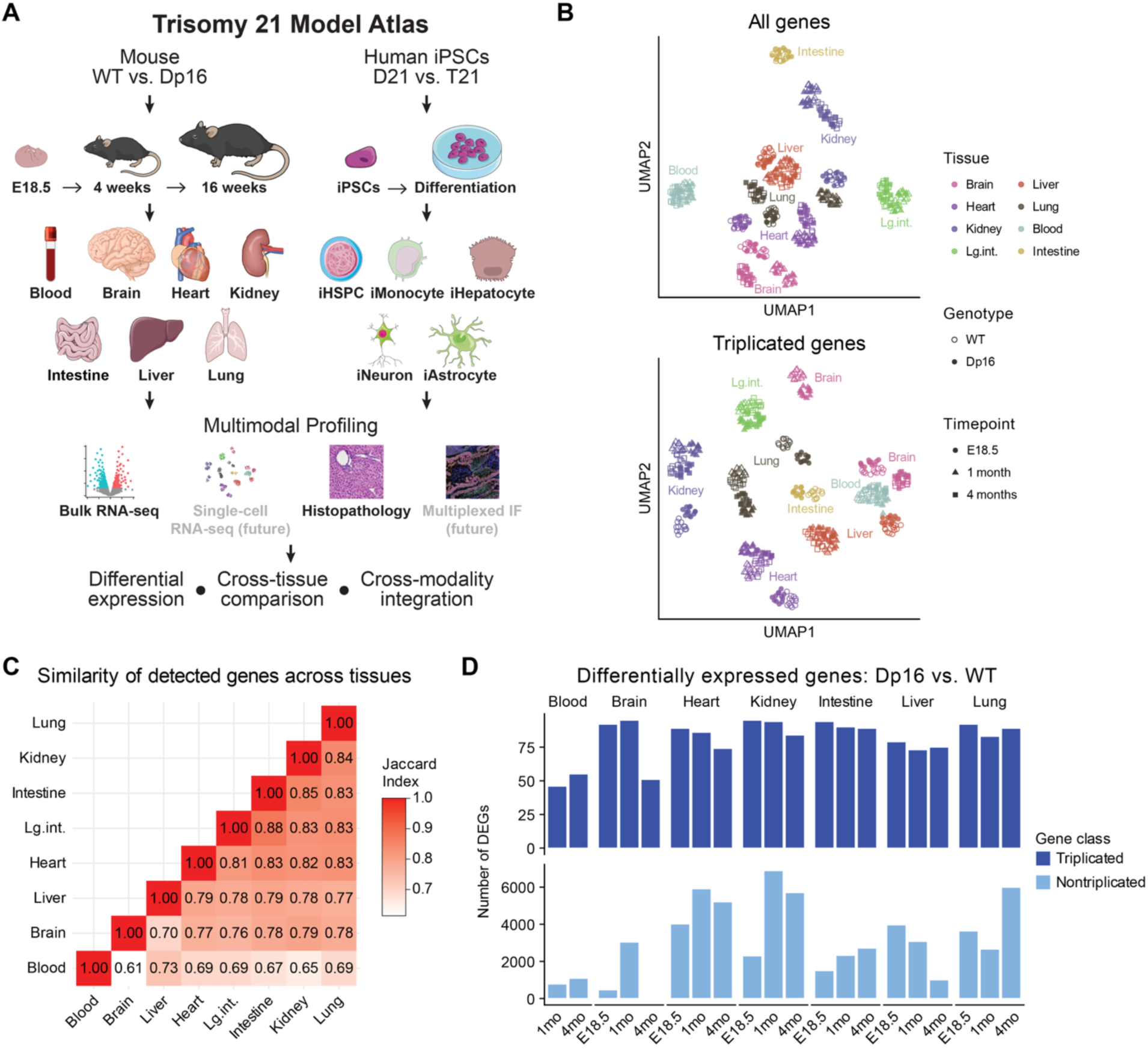
Overview of the Trisomy 21 Model Atlas study design. **(A)** Wild-type (WT) and Dp16 mice were profiled across three developmental timepoints (E18.5, 1 month, 4 months) in seven tissues (blood, brain, heart, kidney, intestine, liver, and lung; 12 WT and 12 Dp16 mice per tissue-timepoint). In parallel, disomic (D21) and trisomic (T21) human iPSCs were differentiated into five cell types (hematopoietic stem/progenitor cells [HSPCs], monocytes, hepatocytes, neurons, and astrocytes; 6 D21 and 6 T21 per differentiation). Mouse tissues and iPSC-derived cell types were profiled by bulk RNA-sequencing and, for mouse tissues, histopathology; multiplexed immunofluorescence and single-cell RNA-sequencing (red) are planned additions to future Atlas releases. Together, these datasets support differential expression analysis, cross-tissue comparison, and cross-model/modality integration. **(B)** UMAP projection of bulk RNA-sequencing data from all seven mouse tissues, computed using all genes (top) or triplicated genes only (bottom). Each point represents one sample, with tissue represented by color (intestine collected as whole intestine at E18.5 and as large intestine at 1 and 4 months), developmental timepoint by shape, and genotype by open (WT) or closed (Dp16) symbols. **(C)** Pairwise Jaccard similarity of detected genes between tissues. **(D)** Global summary of differential expression across the Atlas. Number of differentially expressed triplicated (top, dark blue) and nontriplicated (bottom, light blue) genes (q<0.1) in each tissue at each developmental timepoint; note the different y-axis scales in each panel.

To visualize the transcriptional landscape across the mouse arm of the Atlas, we performed UMAP dimensionality reduction on all tissues, timepoints, and genotypes together. With all expressed genes included, samples separated primarily by tissue (**Figure 1B, top**), consistent with tissue identity as the dominant axis of variation across the dataset. Restricting the analysis to triplicated genes produced greater separation within tissue clusters, in part reflecting genotype-associated substructure that is largely absent when all genes are considered (**Figure 1B, bottom**). This is consistent with a dosage effect acting across all tissues and timepoints but secondary to tissue identity in the overall transcriptional landscape. We note that sequencing batch is partially confounded with tissue and timepoint for a subset of samples, and we therefore do not draw strong biological conclusions from the fine structure of these projections.

Gene detection was broadly consistent across the Atlas, with pairwise Jaccard similarities of detected genes between tissues generally falling between 0.70 and 0.85 (**Figure 1C**), indicating that variation in differentially expressed gene (DEG) number between tissues are not simply a consequence of differing transcriptome coverage. Nonetheless, a substantial fraction of genes were detected in only a subset of tissues, reflecting genuine tissue-restricted expression – most evident for blood, which showed the lowest similarity to other tissues. This combination of broadly shared detection with meaningful tissue-specific expression informed our approach to differential expression testing: a single model spanning all tissues and timepoints would apply uniform low-expression prefiltering across contexts in which a gene may be robustly expressed in one tissue and absent in another, compromising variance estimation for tissue-restricted genes. We therefore performed differential expression testing separately for each tissue-timepoint combination (**Table S2**), an approach that also limits the influence of batch structure on genotype comparisons within a given tissue and timepoint. A higher-resolution view of detected-gene similarity at the tissue-timepoint level (**Figure S1**) confirms that this tissue specificity persists after accounting for developmental stage.

Summarizing differential expression across the Atlas revealed two key global features (**Figure 1D**). First, the majority of triplicated genes are differentially expressed in nearly every tissue-timepoint combination examined, confirming dosage sensitivity as a broadly consistent consequence of triplication. Second, although triplicated genes represent only a small fraction of the genome, their effects propagate into a far broader differential expression landscape that is numerically dominated by nontriplicated genes, which exceed triplicated DEGs by a median of 32-fold (IQR 19–55-fold; range 1– 73-fold) across tissue-timepoints. The contrast is therefore twofold: dosage effects are near-complete in proportional terms but limited in scope, whereas downstream responses are overwhelming in absolute number and highly context-dependent. This asymmetry is easily overlooked when attention centers on chromosome 21 itself, yet it indicates that most of the downstream transcriptional consequences of the trisomy manifest beyond the triplicated region. Given the sensitivity of DEG counts to expression level, sample size, and the batch structure noted above, we treat these summary statistics as an orienting observation; the underlying patterns are examined directly at the level of individual genes and tissues in the sections that follow.

### Conserved gene dosage effects and tissue-specific transcriptional responses

Having established the overall structure of the Atlas, we next examined the transcriptional consequences of trisomy in detail within individual tissues at the 4-month timepoint, where all seven tissues are represented (**Figure S2A**).

The cross-tissue breadth with which genes respond to trisomy differs sharply between triplicated and nontriplicated genes (**Figure 2A**). Every triplicated gene was differentially expressed in at least one tissue, and most were differentially expressed in the majority of tissues, with the largest single group significant in all seven. Nontriplicated genes show the opposite distribution: the vast majority are differentially expressed in only one or two tissues, with progressively fewer genes shared across three or more. Dosage-driven responses are therefore largely tissue-agnostic, while downstream responses are predominantly tissue-restricted.

**Figure 2.**
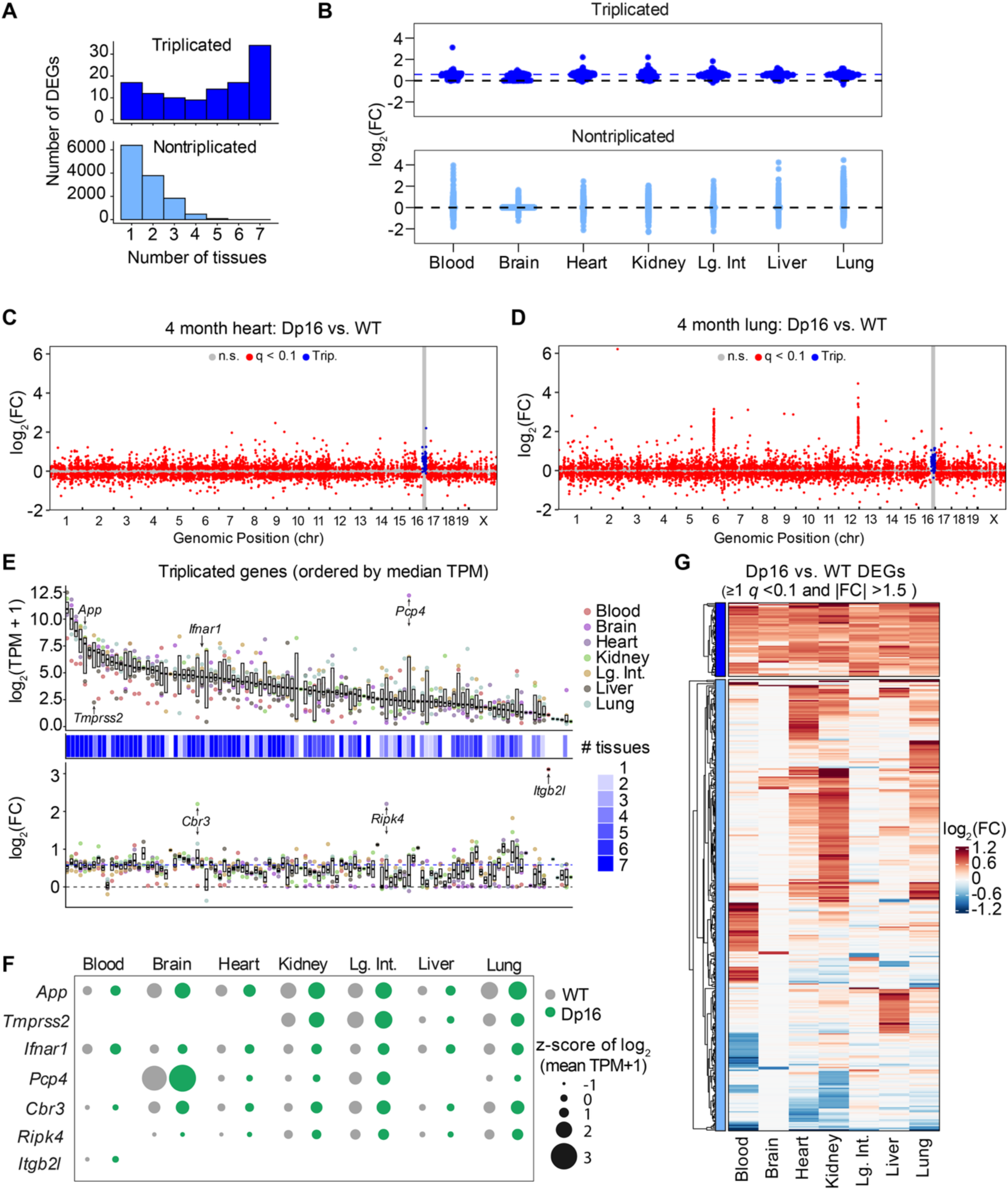
Conserved gene dosage effects and tissue-specific transcriptional responses at the 4-month timepoint. **(A)** Number of triplicated (top, dark blue) and nontriplicated (bottom, light blue) genes differentially expressed (q<0.1) in the indicated number of tissues at 4 months (12 WT and 12 Dp16 mice per tissue); note the different y-axis scales between sub-panels. Genes not differentially expressed in any tissue are not shown. **(B)** Distribution of log_2_ fold-changes (Dp16 vs. WT) for triplicated (top) and nontriplicated (bottom) differentially expressed genes in each tissue at 4 months. **(C, D)** Differentially expressed genes in heart (C) and lung (D) at 4 months by genomic position; the triplicated MMU16 region is shaded in grey. Triplicated genes are shown in blue, nontriplicated genes meeting the significance threshold (q<0.1) in red, and genes not meeting the threshold in grey. **(E)** Triplicated genes ordered by median expression (transcripts per million, TPM) across 4-month tissues. Top, log_2_-transformed TPM per tissue in Dp16 mice. Middle, color bars indicate the number of tissues in which a gene is differentially expressed. Bottom, log_2_-transformed fold-change per tissue. Boxes indicate interquartile ranges across tissues and black bars represent medians. TPM values are adjusted for SVA-derived surrogate variables. **(F)** z-score of log_2_(mean SVA-adjusted TPM + 1) for selected triplicated genes across 4-month tissues in WT (grey) and Dp16 (green) mice; point size increases with z-score. **(G)** Heatmap of log_2_ fold-changes for triplicated and nontriplicated genes across 4-month tissues. Genes with q<0.1 and an absolute fold-change of at least 1.5 in at least one tissue are shown.

A similar asymmetry holds for the magnitude and direction of change (**Figure 2B**). Triplicated genes are uniformly upregulated in every tissue, with fold-changes distributed around the 1.5-fold expectation for a 3:2 gene dosage ratio; formal testing against a null of 1.5-fold in heart, kidney, liver, and lung confirmed that most triplicated genes do not depart significantly from this expectation. Nontriplicated genes, by contrast, are perturbed in both directions and span a considerably wider range of fold-changes in every tissue examined. This distinction matters for interpretation: elevated expression of triplicated genes follows directly and predictably from their copy number, whereas downstream responses vary in both direction and magnitude across tissues, indicating that context strongly shapes the transcriptional landscape of T21.

Plotting by genomic position further illustrates the asymmetry (**Figure 2C, D**). For example, in both heart and lung, the triplicated MMU16 segment appears as a contiguous block of coordinate upregulation, as expected from copy number. Critically, this region represents only a small minority of the differentially expressed genes in either tissue: the great majority lie outside the triplicated region, distributed across all autosomes and the X chromosome with no comparable positional clustering. Although heart and lung produce broadly similar genome-wide patterns at this resolution, the specific genes involved differ substantially between them.

Despite the relatively tight clustering of triplicated DEGs around 1.5-fold upregulation, there is nonetheless variation around this trend. Thus, we next asked whether this variation relates to baseline expression level. Ordering triplicated genes by median expression across tissues reveals no systematic relationship between expression level and fold-change (**Figure 2E**). Fold-changes are tightly clustered near 1.5-fold among highly expressed genes and become progressively more variable as median expression decreases, however, this widening is what would be expected from greater measurement variance at low expression, compounded by the fact that genes expressed at low median levels are often expressed in only a subset of tissues. We therefore find no evidence that dosage proportionality itself weakens with expression level. Tissue breadth, by contrast, does track median expression: triplicated genes expressed at high levels across tissues are differentially expressed in the greatest number of tissues.

Within this overall context, individual triplicated genes illustrate distinct scenarios (**Figure 2E, F**). First, the breadth of the dosage response is bounded by where a gene is normally expressed. *App* and *Ifnar1* are expressed and upregulated consistently across all seven tissues, as expected for genes with broad tissue distribution; *Tmprss2* is abundant in kidney, large intestine, and lung, weakly expressed in liver, and undetected in blood, brain, and heart; *Pcp4* is very highly expressed in brain and only weakly elsewhere. Second, a small number of triplicated genes show fold-changes well above the dosage expectation, including *Cbr3* in kidney and *Ripk4* in heart (∼4.6-fold), and *Itgb2l*, expressed ∼8.7-fold higher in Dp16 than WT blood but undetected in either genotype in any other tissue. Thus, while copy number predicts a uniform 1.5-fold increase, tissue-specific expression and regulation can push individual genes well beyond that expectation – most visibly upward, since suppression below the dosage expectation is harder to resolve against an already-elevated baseline.

Direct comparison of DEGs across tissues confirms that the breadth difference in Figure 2A reflects genuinely distinct gene sets rather than simply differing numbers of genes reaching significance. Hierarchical clustering of fold-changes across tissues shows this directly: triplicated genes form a coherent block of shared upregulation, while nontriplicated genes resolve into tissue-associated groups with minimal cross-tissue conservation (**Figure 2G**). Pairwise Jaccard indices for differentially expressed triplicated genes are uniformly high (0.62–0.97), indicating that largely the same set of triplicated genes responds in every tissue. For nontriplicated genes the corresponding values are much lower (0.02–0.56), with the most divergent comparisons – brain versus blood and brain versus liver – sharing almost no DEGs (**Figure S2B**). Spearman correlations of per-gene fold-changes show the same pattern for triplicated genes (rho 0.38–0.90), indicating that the magnitude of the response, not just the identity of the responding genes, is largely preserved across tissues (**Figure S2C**). Correlation values for nontriplicated genes should be interpreted with care, as the tissue pairs sharing the fewest DEGs yield correlations computed over very few genes and are correspondingly unstable, and can appear spuriously high.

Taken together, these analyses distinguish two components of the transcriptional response to trisomy 21. The primary effect of gene dosage is remarkably uniform – a consistent, near-proportional upregulation of triplicated genes, largely independent of tissue context. The secondary response, which accounts for the overwhelming majority of DEGs, instead differs substantially between tissues in terms of the identity, direction, and magnitude of changes, producing a distinct transcriptional signature in each. Understanding how trisomy 21 produces its diverse phenotypic consequences therefore requires characterizing not only which genes are triplicated, but how each tissue responds to their elevated expression.

### Inflammatory pathways are broadly enriched but driven by tissue-specific gene-level contributions

The tissue-restricted nature of the downstream response raises an apparent paradox: if a largely distinct set of genes is dysregulated in each tissue, how does trisomy 21 produce the recurrent phenotypic themes observed across organ systems in Down syndrome, such as altered growth and immune dysfunction? We therefore asked whether these tissue-specific gene sets nonetheless converge on shared biological programs.

Gene set enrichment analysis across the seven 4-month tissues addresses this directly (**Figure 3A** and **S3A, Table S3**). Several inflammation-related gene sets, including IFNα and IFNγ responses and interleukin/JAK/STAT signaling, are positively enriched across most tissues (**Figure 3A**), consistent with chronic baseline interferon activation as a pervasive feature of the trisomic transcriptome rather than a property of any single tissue or organ. Not all inflammatory gene sets behave uniformly: enrichment of TNFα signaling via NFκB is significantly elevated in lung, heart, and kidney, significantly reduced in brain and blood, and unchanged in large intestine and liver. More importantly, shared enrichment does not by itself indicate a shared underlying response. Enrichment plots for blood and heart show that comparable IFNγ response enrichment can arise from appreciably different gene rankings (**Figure 3B**), prompting us to examine which genes drive the signature in each tissue.

**Figure 3:**
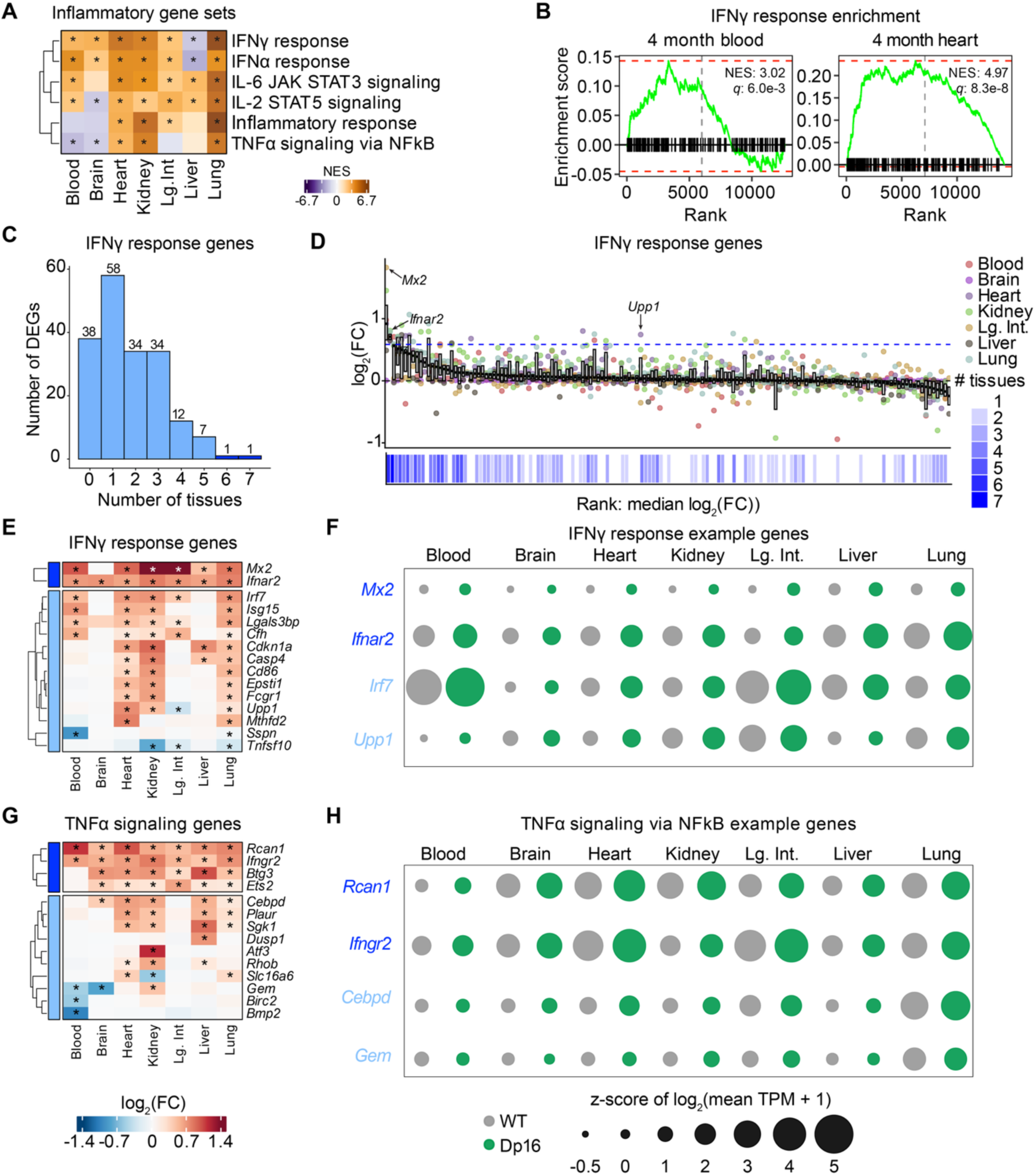
Broad enrichment of inflammatory pathways is driven by tissue-specific gene-level contributions at the 4-month timepoint. **(A)** Heatmap of normalized enrichment scores (NES) for inflammatory gene sets defined by Gene Set Enrichment Analysis (GSEA) across 4-month tissues (12 WT and 12 Dp16 mice per tissue). Asterisks indicate q<0.1. **(B)** Enrichment plots for the IFNγ response gene set in blood and heart at 4 months. **(C)** Number of IFNγ response genes differentially expressed (q<0.1) in the indicated number of tissues at 4 months. **(D)** IFNγ response genes ranked by median log_2_ fold-change across 4-month tissues; each point represents one gene in one tissue, colored by tissue. The blue color bar below the main plot indicates the number of tissues in which each gene is differentially expressed. **(E)** Heatmap of log_2_ fold-changes for triplicated and nontriplicated IFNγ response genes across 4-month tissues. Genes with q<0.1 in any tissue and an absolute fold-change of at least 1.5 are shown. **(F)** z-score of log_2_(mean SVA-adjusted TPM + 1) for selected IFNγ response genes across 4-month tissues in WT (grey) and Dp16 (green) mice; point size increases with z-score. **(G)** As in (E), for TNFα signaling via NFκB gene set members. **(H)** As in (F), for selected TNFα signaling via NFκB gene set members.

Examination of which genes contributing to the shared interferon signature revealed that convergence at the pathway level does not guarantee convergence at the gene level (**Figure 3C**). Binning IFNγ-related genes by the number of tissues in which they are differentially expressed produces a distribution closely resembling that of nontriplicated genes genome-wide: a plurality are differentially expressed in only one tissue, with progressively fewer shared across more. *Ifnar2* and *Mx2*, the two genes triplicated in Dp16, are the conspicuous exception, differentially expressed in six or seven tissues. This single gene set thus recapitulates the broader architecture of the trisomic transcriptome: a small set of dosage-driven genes responding almost everywhere, combined with a much larger set of tissue-restricted genes that converge on a common pathway-level signature.

Fold-change magnitude reinforces this division (**Figure 3D-F**). Ranking IFNγ-related genes by median fold-change across tissues shows that even the triplicated genes are not perfectly uniform: *Mx2* shows larger fold-changes in kidney and large intestine than elsewhere, whereas *Ifnar2* responds to a similar degree in every tissue. Nontriplicated genes diverge considerably more. *Irf7* is consistently upregulated in heart, kidney, large intestine, lung, and blood but unchanged in brain and liver, and its baseline expression varies widely (high in blood and large intestine, low elsewhere). *Upp1* adds a further layer of complexity: it is upregulated in heart, kidney, and lung but downregulated in large intestine. A gene may therefore contribute to interferon enrichment in one tissue while moving in the opposite direction in another.

TNFα-related genes display the same general pattern, albeit with even greater tissue divergence (**Figure 3G-H**). The triplicated genes *Rcan1* and *Ifngr2* are consistently upregulated across tissues, differing mainly in magnitude. Among nontriplicated genes, *Cebpd* is upregulated in five of seven tissues but is abundantly expressed only in lung, while *Gem* is upregulated in kidney, downregulated in blood and brain, and unchanged in the remaining four. These are not subtle quantitative differences around a common response: the same annotated gene set may contain genes changing in opposite directions across organs, indicating that a shared pathway-level enrichment score can mask substantial heterogeneity in the underlying transcriptional response. Pathway-level enrichment statistics alone cannot distinguish these programs from one another; only inspection of the underlying genes reveals how each contributes in a given context.

Taken together, these analyses show how a shared molecular hallmark of trisomy 21 can coexist with highly tissue-specific gene expression. Chronic IFN activation is one of the most consistently reported features of Down syndrome across human cohorts and mouse models^24,28–35^, and the Atlas datasets corroborates this as a pathway-level signature detectable in nearly every tissue examined. However, that signature is composed of substantially different genes in each tissue. Pathway-level agreement across tissues should therefore not be taken to imply a common underlying transcriptional program – a distinction with direct consequences for interpreting cross-tissue and cross-model comparisons, and for anticipating which tissues will respond similarly to a pathway-directed intervention.

### Developmental context modulates downstream transcriptional responses to trisomy 21

Having established that tissue identity is a primary organizing principle of the transcriptional response to trisomy at a single developmental stage, we next asked whether this architecture carries across development, or whether the balance between conserved dosage effects and tissue-specific downstream responses shifts as tissues mature.

Comparing fold-changes across E18.5, 1-month, and 4-month timepoints shows that the core asymmetry identified at 4 months holds across each developmental timepoint examined (**Figure 4A, Table S2**). Triplicated genes are consistently upregulated in Dp16 relative to WT at all three timepoints, whereas nontriplicated genes are upregulated, downregulated, or unchanged depending on the specific tissue and timepoint. The same asymmetry holds in breadth: at every timepoint, triplicated genes are typically differentially expressed in six or seven tissues, while nontriplicated genes are restricted to one or two (**Figure 4B**). The tissue-agnostic dosage effect and the tissue-restricted nature of the downstream response are therefore not features specific to the 4-month timepoint characterized above, but persist from late embryogenesis through early adulthood.

**Figure 4.**
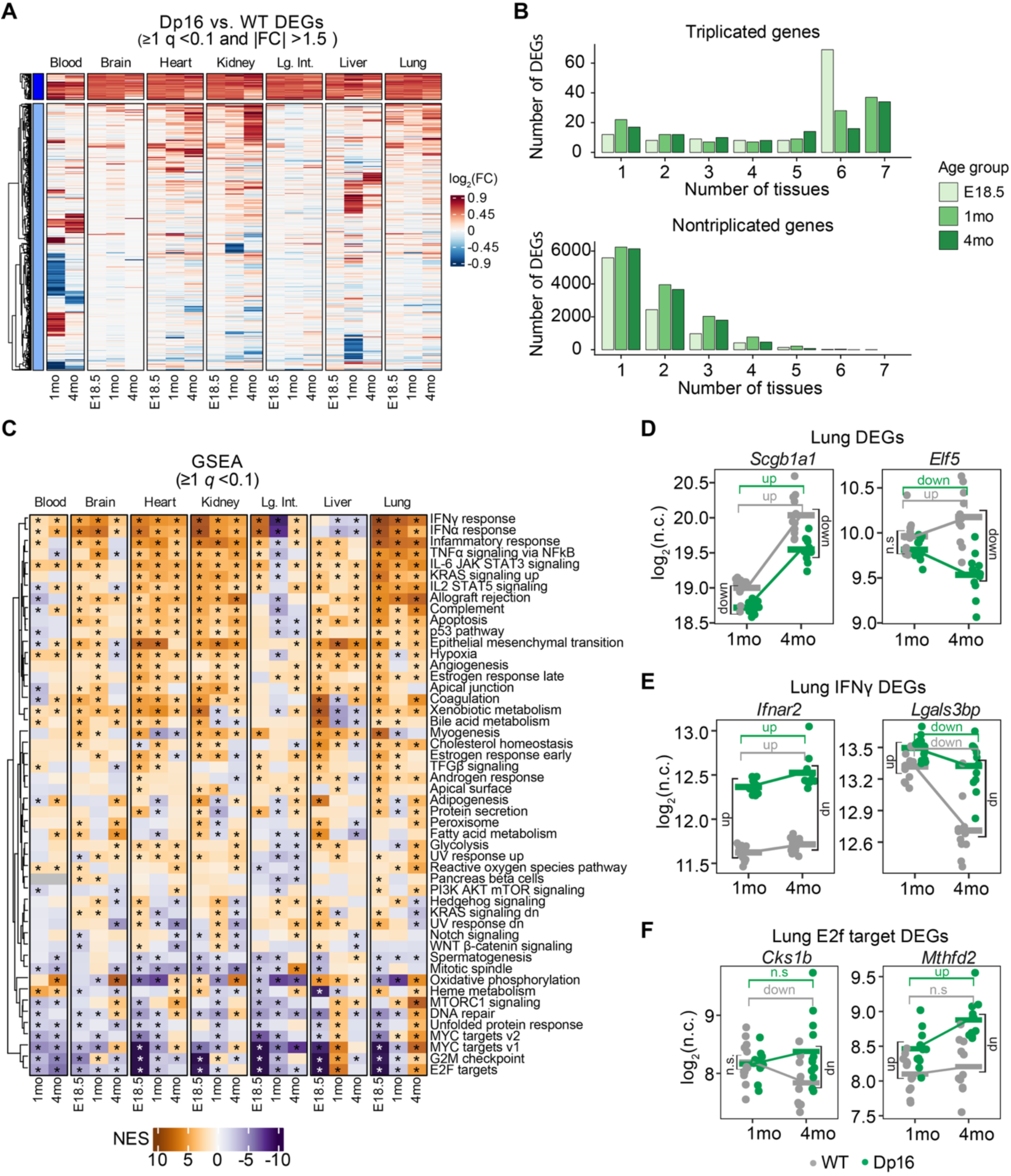
Developmental context shapes downstream transcriptional responses to trisomy 21. **(A)** Heatmap of log_2_ fold-changes for triplicated and nontriplicated genes across E18.5, 1-month, and 4-month timepoints in all tissues (12 WT and 12 Dp16 mice per tissue-timepoint). Genes with q<0.1 in any tissue-timepoint and an absolute fold-change of at least 1.5 are shown. **(B)** Number of tissues in which triplicated (top) and nontriplicated (bottom) genes are differentially expressed (q<0.1), shown separately for E18.5, 1-month, and 4-month timepoints. **(C)** Heatmap of normalized enrichment scores (NES) from Gene Set Enrichment Analysis (GSEA) of Hallmark gene sets across E18.5, 1-month, and 4-month timepoints in each tissue. Pathways with q<0.1 in at least one tissue-timepoint are shown. Asterisks indicate q<0.1. **(D)** Relative expression of example genes in lung (*Scgb1a1*, *Elf5*) from WT and Dp16 mice at 1 and 4 months. Log_2_-transformed normalized counts are shown. **(E)** As in (D), for example IFNγ response genes (*Ifnar2*, *Lgals3bp*). **(F)** As in (D), for example E2F target genes (*Cks1b*, *Mthfd2*).

As an example, examining the lung across timepoints in more detail reveals a more nuanced picture (**Figure S4A, B**). The specific triplicated genes reaching significance are nearly identical at every timepoint (Jaccard 0.95–0.98), yet the magnitude of their fold-changes is comparatively more variable across timepoints (rho 0.51–0.69). However, this may partly reflect the narrow range of triplicated gene fold-changes clustered near the 1.5-fold dosage expectation, which limits the resolution of rank-based correlation. Nontriplicated genes show the inverse relationship: which specific genes reach significance changes considerably between timepoints (Jaccard 0.39–0.42), but among DEGs shared across more than one timepoint, the magnitude of change is relatively well conserved (rho 0.63–0.85). In other words, the dosage response involves a relatively stable set of changes whose individual effect sizes drift somewhat with age, while the downstream response involves differing genes whose magnitude, once engaged, is comparatively consistent.

At the pathway level, a similar pattern of time-invariant and time-dependent responses is apparent (**Figure 4C, Table S3 and Table S4**). Interferon-related gene sets, including IFNγ and IFNα response, are enriched in Dp16 relative to WT across nearly every tissue and timepoint, indicating that the persistent inflammatory signature described above is already present at the earliest stage examined. Cell-cycle-related gene sets behave very differently: E2F targets and G2M checkpoint signatures are strongly depleted in Dp16 at E18.5 in heart, kidney, large intestine, liver, and lung, but this depletion is markedly reduced or reversed by later timepoints. Inflammatory signaling thus appears to be an early and durable consequence of the trisomy, whereas cell-cycle pathways are perturbed specifically during a defined developmental window, possibly reflecting the transient proliferative demands of late embryonic tissue growth.

Individual genes in the lung illustrate the range of trajectories underlying these pathway-level patterns (**Figure 4D–F**). Some differences are established early and persist: *Scgb1a1*, a marker of airway club cells, is expressed at lower levels in Dp16 vs. WT at 1 month, and this relative deficit is maintained at 4 months even as expression increases in both genotypes (**Figure 4D, Table S5**). Others emerge only with maturation: *Elf5* does not differ between genotypes at 1 month, but by 4 months WT expression has increased while Dp16 expression has decreased, producing a difference where none previously existed. Among IFNγ-response genes, *Ifnar2* follows the pattern expected of a dosage-driven gene, exceeding WT expression at both timepoints and increasing further with age, while *Lgals3bp* instead narrows over time as WT expression declines more steeply than Dp16 between 1 and 4 months (**Figure 4E**). Among E2F target genes, *Cks1b* becomes significantly different only at 4 months, driven by a decline in WT rather than a change in Dp16, while *Mthfd2* shows the opposite trajectory, rising progressively in Dp16 while WT remains flat (**Figure 4F**). Persistent, emergent, converging, and diverging trajectories are therefore all represented within a single tissue, underscoring the idea that developmental timing does not act along a single axis but produces genuinely distinct dynamic relationships between genotypes at individual genes.

Taken together, these analyses show that the core architecture of the trisomic transcriptome is a near-universal, dosage-proportional upregulation of triplicated genes against a much larger and more variable nontriplicated response that is established by late embryogenesis and maintained through early adulthood. Superimposed on this stable architecture is the developmental dimension: the magnitude of triplicated gene responses shifts somewhat with age even as the responding genes remain the same, cell-cycle pathways are altered in a developmentally restricted window largely absent from the later timepoints examined, and individual downstream genes can gain, lose, or reverse a genotype difference as tissues mature. Fully resolving these dynamics, including whether they generalize beyond the lung and heart, kidney, large intestine, and liver examples shown here will require deeper analysis alongside additional developmental sampling, tissues, and cell types planned for future Atlas releases.

### Sex modifies tissue-specific transcriptional responses to trisomy 21

Building on the tissue- and stage-dependent architecture described above, we next examined whether the transcriptional response to trisomy differs by sex. This comparison is inherently underpowered relative to the genotype comparisons presented above. Splitting each tissue by sex reduces the number of animals per group to six, so these results should be read as indicating sex-dependent patterns worth further investigation rather than as a definitive characterization.

Comparing trisomy-associated fold-changes in male and female animals directly shows that triplicated genes behave consistently between the sexes in most tissues (**Figure 5A; Figure S5A, Table S6**). For example, in heart and lung, differentially expressed triplicated genes overlap almost completely between sexes (Jaccard = 0.92 and 0.96, respectively), extending, with somewhat lower overlap, to kidney (0.89). Liver is the clear exception, with the Jaccard index falling to 0.38, driven in part by a larger set of triplicated genes reaching significance in males than in females. *Hspa13* and *Morc3* illustrate the more typical pattern, upregulated by comparable magnitudes in both sexes in heart/lung and kidney/large intestine respectively, while *Wrb* illustrates the exception, upregulated in both sexes in blood but only in males in liver (**Figure 5B**). Even where the same triplicated genes are differentially expressed in both sexes, rank correlations of their fold-changes between sexes are modest and variable across tissues (rho -0.11 to 0.68; **Figure S5A**). However, this may partly reflect the narrow range of triplicated gene fold-changes clustered near the 1.5-fold dosage expectation, which limits the resolution of rank-based correlation.

**Figure 5.**
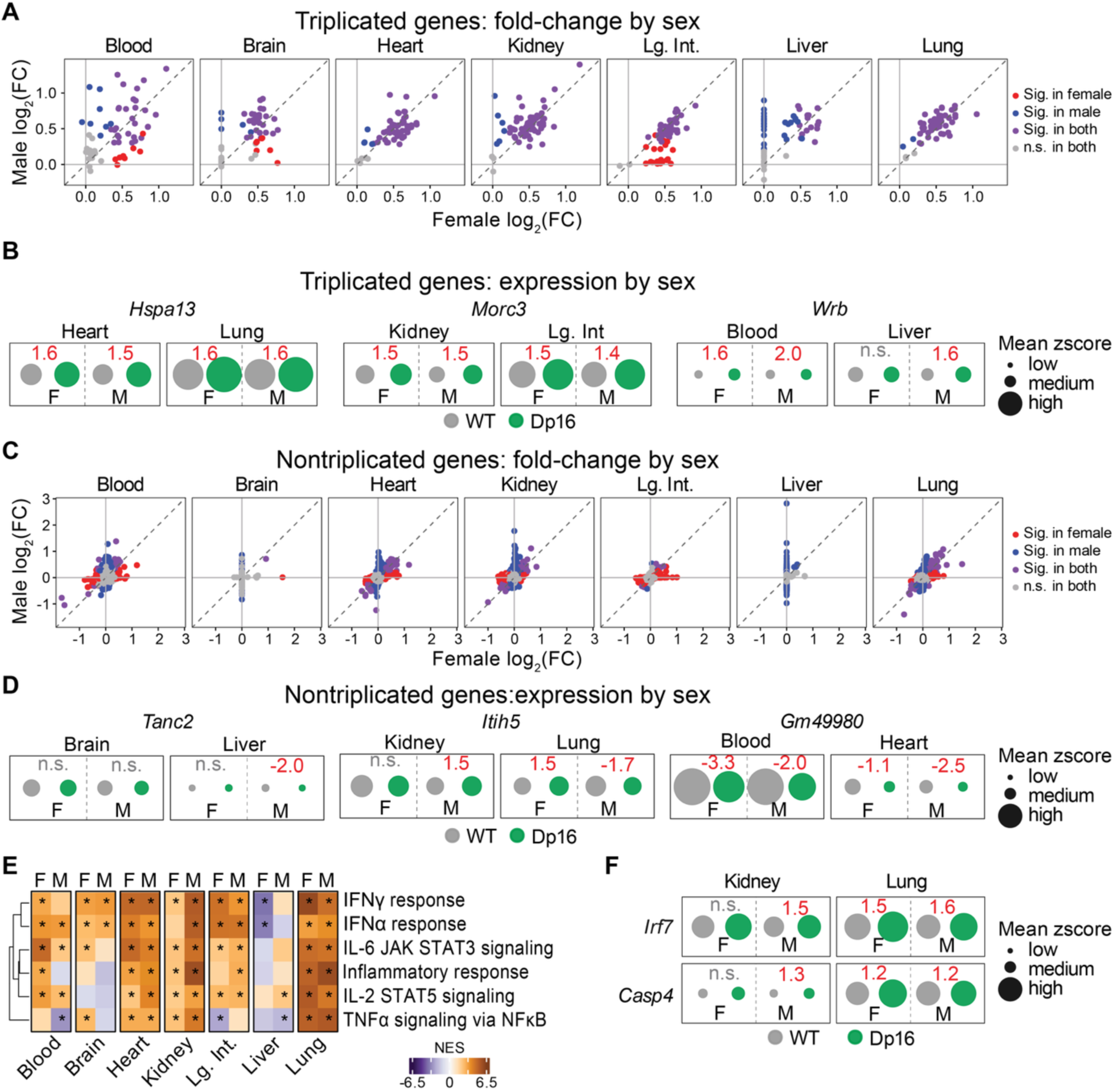
Sex modifies tissue-specific transcriptional responses to trisomy 21 at 4 months. **(A)** Scatterplot of log_2_ fold-change (Dp16 vs. WT) for triplicated genes in females (x-axis) and males (y-axis) across seven tissues (6 WT and 6 Dp16 mice per sex-timepoint-tissue group). DEGs (q<0.1) in females only are shown in red, males only in blue, both sexes in purple, and neither sex in grey. **(B)** Mean z-score of SVA-adjusted TPM for selected triplicated genes (*Hspa13*, *Morc3*, *Wrb*) in WT and Dp16 females (F) and males (M) in the indicated tissues; point size increases with z-score. Fold-change values are shown in red where significant (q<0.1) and grey (n.s.) where not. **(C)** As in (A), for nontriplicated genes. **(D)** As in (B), for selected nontriplicated genes (*Tanc2*, *Itih5*, *Gm49980*). **(E)** Heatmap of normalized enrichment scores (NES) for inflammatory gene sets defined by Gene Set Enrichment Analysis (GSEA), shown separately for females (F) and males (M) in each tissue. Asterisks indicate q<0.1. **(F)** As in (B), for selected IFNγ response genes (*Irf7*, *Casp4*) in kidney and lung.

Nontriplicated genes show substantially lower overlap between sexes across every tissue examined (**Figure 5C; Figure S5B**). Jaccard indices for nontriplicated DEGs range from 0.00 to 0.14, an order of magnitude below the corresponding triplicated values. This low overlap should be interpreted cautiously, as genes near the significance threshold in one sex may fail to reach it in the other for reasons unrelated to sex. Consistent with this, fold-changes for genes reaching significance in both sexes are moderately correlated (rho 0.12–0.65), indicating that the underlying transcriptional responses are more similar than the overlap statistics alone would suggest. Nonetheless individual example genes illustrate patterns that may be worth further examination: *Tanc2* is unchanged in both sexes in brain but downregulated in the male liver, while *Itih5* is upregulated in male kidney and shows discordant directions in the lung (**Figure 5D**). Others appear robust to sex; *Gm49980*, for example, is downregulated in both sexes in blood and heart. Distinguishing genuine sex-dependent responses from threshold effects will require larger cohorts than the present analysis affords.

At the pathway level, inflammatory-related gene sets are again broadly enriched in both sexes across most tissues, with occasional sex-dependent differences in degree (**Figure 5E, Table S7**). In lung, IFNγ, IFNα, IL-6/JAK/STAT3, inflammatory response, IL-2/STAT5, and TNFα signaling via NFκB are all enriched in both females and males. In kidney, the same gene sets, with the exception of IFNα, are enriched in both sexes, but enrichment scores are consistently lower in females than males. This is illustrated at the gene level by two IFNγ-response genes, *Irf7* and *Casp4*, both significantly upregulated in females and males in lung but reaching significance only in males in kidney (**Figure 5F**). However, as above, this asymmetry may in part reflect reduced power in the female subgroup rather than a true absence of effect. As with the tissue-specific pathway convergence described in Figure 3, shared pathway-level enrichment between sexes does not guarantee a shared magnitude of response at the level of individual genes.

Taken together, these analyses indicate that sex has a comparatively modest influence on the core dosage signature. Most triplicated genes respond similarly in males and females, while nontriplicated genes show substantially lower concordance between sexes, a pattern that may reflect genuine sex-dependent regulation, reduced statistical power, or both. Sex thus joins developmental stage as a further axis of biological context worth accounting for in the transcriptional response to trisomy within a given tissue. Given the reduced power inherent to this sex-stratified analysis, these patterns are best treated as a preliminary map for hypothesis generation, to be refined as sample sizes grow in future Atlas releases and worth prioritizing for further investigation given their potential relevance to sex differences in Down syndrome phenotypes.

### Integration of histopathology and transcriptomics links gene expression to tissue pathology

To connect transcriptional changes characterized thus far with tissue-level physiology, we performed blinded histopathological grading on tissues from the same animals used for transcriptomic profiling At E18.5, ten organ systems were evaluated by mid-sagittal whole-embryo sectioning, and no significant genotype differences were detected at this stage (**Table S8)**. At 1 month and 4 months (**Table S9 and Table S10, respectively**), twelve individual tissues were scored using tissue-specific component criteria; component scores were summed within each tissue to produce a composite pathology score for comparison between genotypes.

Composite pathology scores differed significantly between genotypes in a small subset of tissue-timepoint combinations (**Figure 6A**): bone marrow at 1 month, and bone marrow, kidney, and lung at 4 months (Wilcoxon test, effect size, p<0.05). Individual animal-level scores for these four comparisons show Dp16 mice consistently trending toward higher composite pathology than WT, with considerable variability between animals (**Figure 6B**). More tissues reach significance by the later timepoint: only one tissue does so at 1 month, three do so at 4 months, and roughly half of the tissues surveyed show no significant composite difference at either timepoint.

**Figure 6.**
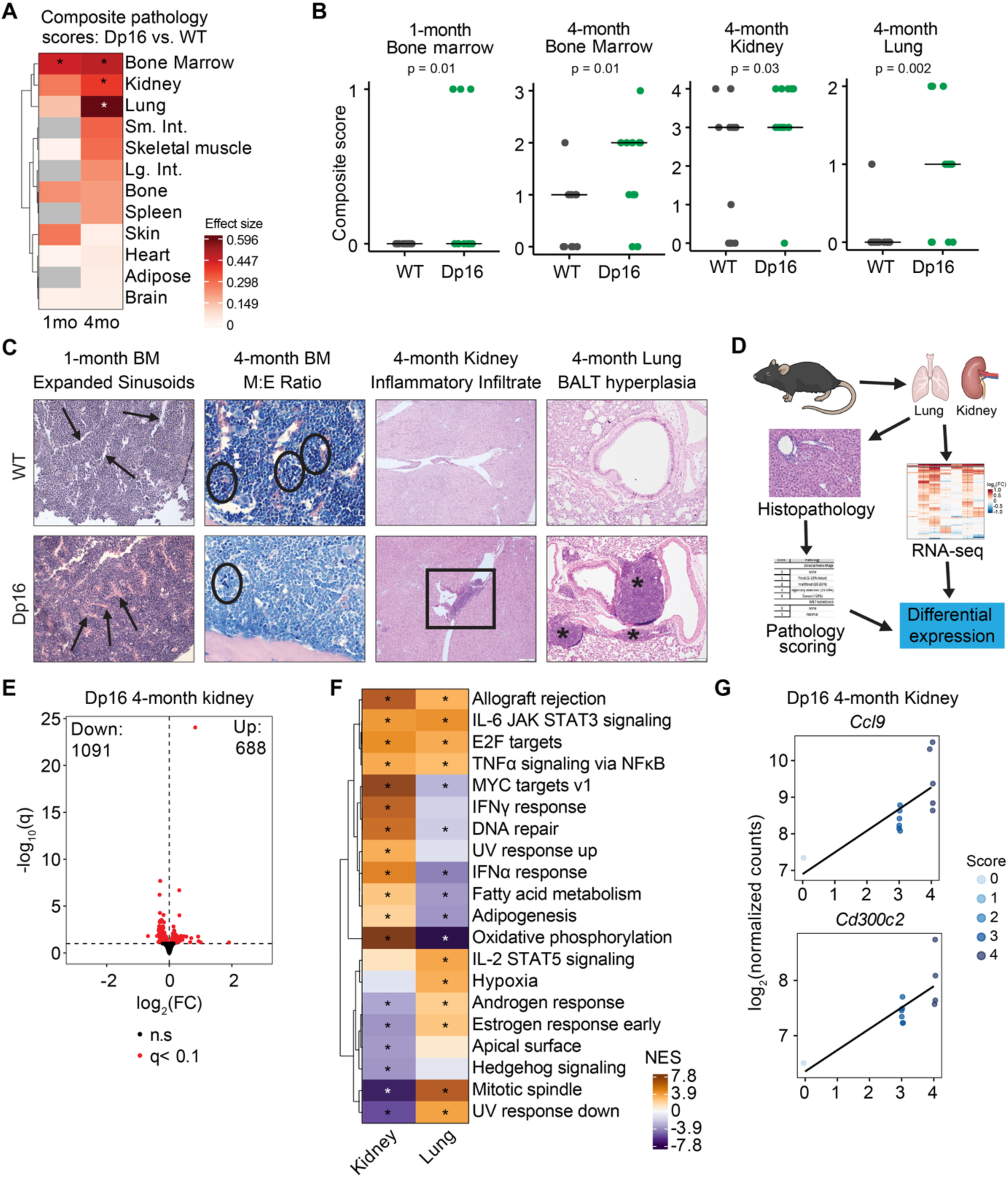
Integration of histopathology and transcriptomics links gene expression to tissue pathology. **(A)** Heatmap of composite histopathology scores (effect size, Dp16 vs. WT) at 1 and 4 months across twelve tissues (12 WT and 12 Dp16 mice per tissue-timepoint). Grey indicates no pathology detected in either genotype (small intestine, colon, spleen) or tissue not collected (adipose). Asterisks indicate a significant difference in effect size (Wilcoxon test, p<0.05). **(B)** Composite pathology scores for the four tissue-timepoint combinations reaching significance in (A). **(C)** Representative histopathological images of WT and Dp16 tissues. Arrows indicate expanded sinusoids/stromal elements (1-month bone marrow); circles indicate erythroid islands (4-month bone marrow); box indicates inflammatory infiltrate (4-month kidney); asterisks indicate lymphocyte and plasma cell aggregates characteristic of BALT hyperplasia (4-month lung). **(D)** Workflow for integrating histopathological grading with transcriptomic profiling in the same animals. **(E)** Volcano plot of genes significantly associated (q<0.1) with composite pathology score in 4-month Dp16 kidney, modeled using DESeq2 with composite score as a continuous covariate. **(F**) Heatmap of normalized enrichment scores (NES) from Gene Set Enrichment Analysis (GSEA) of genes ranked by association with composite pathology score in 4-month Dp16 kidney and lung. Dots indicate q<0.1. **(G)** Expression of example inflammation-related genes (*Ccl9*, *Cd300c2*) as a function of composite pathology score in 4-month Dp16 kidney; point color indicates individual component score.

Representative histology images illustrate the specific pathological features underlying these composite scores (**Figure 6C**). In 1-month bone marrow, Dp16 mice show more prevalent stromal elements, including expanded sinusoids, compared to WT. By 4 months, Dp16 bone marrow shows an increased myeloid-to-erythroid ratio, a key indicator of altered hematopoiesis, with expansion of erythroid islands evident. In 4-month kidney, Dp16 mice display increased inflammatory infiltrates, and in 4-month lung, the dominant finding is bronchial-associated lymphoid tissue (BALT) hyperplasia, characterized by enlarged peribronchial aggregates of lymphocytes and plasma cells. Component-level scoring across all tissues (**Figure S6A**) and quantification of these four specific features (**Figure S6C**) support these qualitative observations.

Enabled by these matched measurements, we next asked whether these pathological features are associated with changes in the transcriptional landscape. To this end, we modeled gene expression in Dp16 kidney and lung as a function of composite pathology score, treated as a continuous covariate within genotype, using DESeq2 (**Figure 6D, Table S11**). Restricting this analysis to Dp16 animals reduces statistical power, so these results are best suited to identifying candidate genes rather than precise effect-size estimation; the more limited range of pathology severity captured by the lung’s single dominant finding, relative to kidney’s broader composite score, likely also contributes to fewer genes reaching significance there. With those caveats, the two tissues nonetheless show a striking difference in scale (**Figure 6E; Figure S6D**): kidney gene expression shows extensive associations with pathology score, with 1,091 genes negatively and 688 positively associated (q<0.1), while lung shows far fewer, with only 8 genes negatively and 13 positively associated.

Gene set enrichment analysis of all genes, ranked by their degree of association with composite pathology score, shows that kidney and lung share a core set of enriched and depleted pathways despite the large difference in the number of individually significant genes (**Figure 6F**). Allograft rejection, IL-6/JAK/STAT3 signaling, E2F targets, and TNFα signaling via NFκB are positively enriched with increasing scores in both tissues, consistent with the broadly inflammatory character of pathology in both organs. Most other pathways instead show tissue-specific, often opposite associations with pathology severity: oxidative phosphorylation, for example, is positively enriched with increasing kidney pathology score but negatively enriched with increasing lung pathology score, while IL-2/STAT5 signaling shows the reverse pattern. The relationship between pathology and transcription therefore resembles the broader architecture of the trisomic transcriptome described earlier in this Atlas. However, the shared core response common to both tissues seen here is comparatively small relative to the extensive tissue-specific divergence.

Individual genes point to possible mechanistic links between pathology and gene expression. In kidney, the inflammation-related genes *Ccl9* and *Cd300c2* rank among the strongest correlates of composite score (**Figure 6G**), consistent with the inflammatory infiltrate observed histologically. In lung, the hypoxia-related genes *Ramp3* and *Arg2* correlate with composite score (**Figure S6E**), suggesting that tissue remodeling in BALT hyperplasia may be accompanied by localized hypoxic signaling.

Taken together, these analyses show that the transcriptional consequences of trisomy 21 are accompanied by measurable tissue pathology in a subset of organs, most prominently bone marrow, kidney, and lung, and that the molecular signature of this pathology recapitulates the broader architecture of the trisomic transcriptome described throughout this Atlas: a shared core response, here centered on inflammatory signaling, alongside a tissue-specific secondary response whose scale tracks the extent of pathology in that organ. This disconnect between the near-universal transcriptional dysregulation caused by trisomy and the more selective, tissue-restricted pattern of histologically detectable pathology suggests that transcriptional change is necessary but not sufficient to produce overt structural change at the timepoints examined here, and that longer-term or more sensitive assessments, planned for future Atlas releases, will be needed to determine whether additional tissues develop pathology at any stage.

### Human iPSC-derived cell types recapitulate conserved and context-specific transcriptional responses to trisomy 21

Finally, having characterized the transcriptional consequences of trisomy 21 across mouse tissues, developmental stages, and sexes, we next asked whether the same core asymmetry of a conserved dosage effect set within a much larger, context-specific downstream response extends to human cells. From a panel of twelve iPSC lines derived from participants in the Human Trisome Project (six trisomic, six disomic for chromosome 21; eight male and four female donors) we carried out differentiations into five distinct cell types: astrocytes, neurons, hematopoietic stem and progenitor cells (HSPCs), monocytes, and hepatocytes (see Methods). Throughout, we use the prefix “i-” to denote iPSC-derived cells (iAstrocytes, iNeurons, iHSPCs, iMonocytes, iHepatocytes), distinguishing them from their primary counterparts.

Differentiation was validated by cell-type-specific markers at both the protein and transcript level (**Figure 7A–C**). For iHSPCs, CD34/CD45 double-positive cells were confirmed by flow cytometry and reported as a frequency of viable cells (**Figure 7A**). For iMonocytes, CD14-positivity was confirmed via flow cytometry and reported as a frequency of viable cells (**Figure 7B)**. Validation of iAstrocytes and iNeurons, along with an extended analysis of the bulk RNA-sequencing data from these two cell types, is described separately^36^; iHepatocyte characterization and sequencing data were reported previously^14^. At the transcript level, expression of the pluripotency markers *POU5F1* and *NANOG* was restricted to undifferentiated iPSCs, while each differentiated cell type expressed canonical identity markers: *GFAP* in iAstrocytes, *MAP2* in iNeurons, *CD34* in iHSPCs, *CD14* in iMonocytes, and *HNF4A* in iHepatocytes (**Figure 7C**).

**Figure 7.**
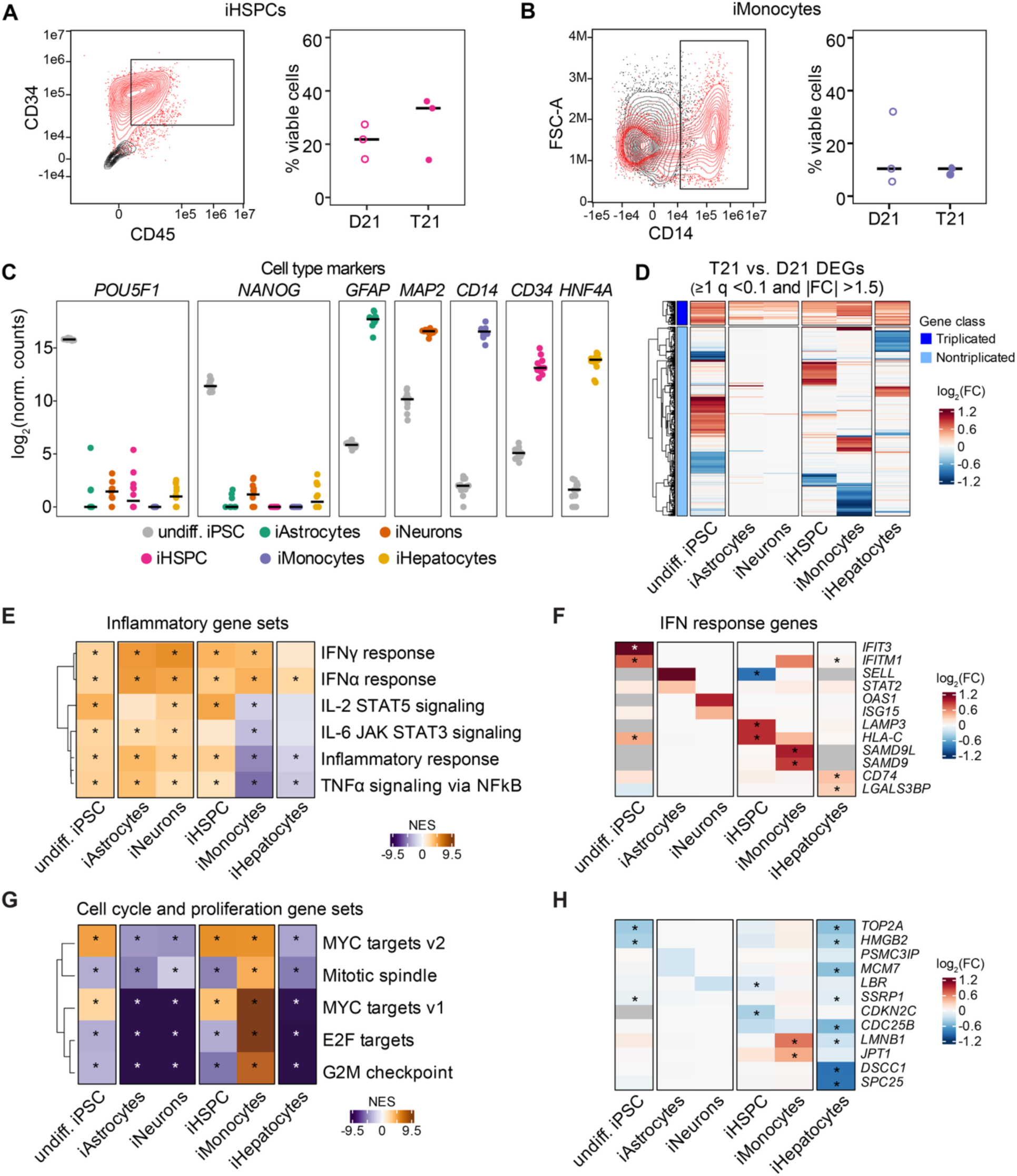
Human iPSC-derived cell types recapitulate conserved dosage effects and cell-type-specific transcriptional responses to trisomy 21. **(A)** Flow cytometry contour plots of iPSC-derived hematopoietic stem and progenitor cells (iHSPCs; CD34 vs. CD45, left) and the percentage of viable cells that are CD34⁺CD45⁺ (right). Red and black contours represent stained and unstained samples, respectively. Gates indicate the population enriched by magnetic separation prior to RNA isolation. Three D21 and three T21 cell lines were measured in triplicate; points represent medians of triplicate measurements and bars represent medians across cell lines. Open points, D21; closed points, T21. **(B)** As in (A), for iMonocytes (FSC-A vs. CD14). **(C)** Expression of pluripotency markers (*POU5F1*, *NANOG*) and cell-type identity markers (*GFAP*, iAstrocytes; *MAP2*, iNeurons; *CD14*, iMonocytes; *CD34*, iHSPCs; *HNF4A*, iHepatocytes) across undifferentiated iPSCs and each differentiated cell type. Plots display log_2_(normalized counts). Cell types comprise 6 D21 and 6 T21 cell lines. **(D)** Heatmap of log_2_ fold-changes (T21 vs. D21) for triplicated and nontriplicated genes across cell types. Genes with q<0.1 in any cell type and an absolute fold-change of at least 1.5 are shown. **(E)** Heatmap of normalized enrichment scores (NES) for inflammatory gene sets defined by Gene Set Enrichment Analysis (GSEA) across cell types. Asterisks indicate q<0.1. **(F)** Heatmap of log_2_ fold-changes for leading-edge genes from the IFNα response gene set across cell types. Top two leading-edge genes per cell type are shown. **(G)** As in (E), for cell-cycle-related gene sets. **(H)** As in (F), for leading-edge nontriplicated genes from the E2F targets gene set.

Comparing differential expression (T21 vs. D21) across these cell types recapitulates the tiered architecture described throughout this Atlas (**Figure 7D and S7A, Table S12**). Triplicated genes are upregulated with substantial overlap across cell types, while nontriplicated genes show comparatively little overlap between them – the same global pattern of tissue-agnostic dosage effects together with context-specific downstream responses observed across mouse tissues, here reproduced across human cell types differentiated from a common panel of donor-derived lines. iAstrocytes and iNeurons stand out as showing fewer differentially expressed genes overall (**Figure S7A**).

At the pathway level, gene set enrichment analysis across all cell types (**Figure S7B, Table S13**) shows inflammatory gene sets – most prominently IFNα and IFNγ response – again enriched across nearly every cell type (**Figure 7E**), consistent with the pervasive interferon signature observed in mouse tissues. Again, as with the mouse signature, this shared enrichment is not built from a fully conserved set of genes: leading-edge genes from the IFNα response gene set differ substantially between cell types (**Figure 7F**), indicating that the generalized inflammatory signature seen in iPSC-derived cells is assembled from distinct cell-type-specific transcriptional programs.

Cell-cycle-related gene sets show a markedly different pattern. Myc targets, mitotic spindle, E2F targets, and G2M checkpoint gene sets are enriched in iMonocytes, variably enriched or depleted in iHSPCs and undifferentiated iPSCs, and strongly depleted in iAstrocytes, iNeurons, and iHepatocytes (**Figure 7G**), with the underlying E2F target leading-edge genes varying correspondingly across cell types (**Figure 7H**). In contrast to the interferon signature, which is enriched in a consistent direction across nearly every cell type, cell-cycle programs are therefore perturbed in opposite directions depending on cellular context.

Taken together, these analyses show that the two-tier architecture of the transcriptional response to trisomy 21 – a conserved dosage effect embedded within a much larger, context-specific downstream response – is not unique to the mouse, but is recapitulated across human iPSC-derived cell types spanning multiple germ layers and lineages. That downstream responses vary with cellular context is expected; what the Atlas provides is an openly available, systematic account of how they vary – which genes and pathways are engaged in which contexts, and which are conserved. This specificity is what mechanistic follow-up and intervention design require, since a therapeutic strategy directed at a pathway shared across tissues may nonetheless act on different genes, and to different effect, in each. A natural extension of this work is a direct, gene-level comparison between matched mouse tissues and human cell types, for example, Dp16 liver against iHepatocytes, A comparison would determine how closely specific downstream responses are conserved across species as well as across cellular contexts; we plan to pursue this cross-model comparison as the Atlas expands beyond this initial data release.

## DISCUSSION

A substantial body of research has linked specific triplicated genes to phenotypes and co-occurring conditions in Down syndrome. Well-characterized examples include APP and early-onset Alzheimer’s disease^37^, HMGN1 and congenital heart defects^38^, and DYRK1A to both congenital heart defects and craniofacial abnormalities^31,39,40^. The interferon receptor gene cluster has been implicated most broadly, contributing to congenital heart defects, craniofacial abnormalities, and immune dysregulation^24^. This approach has been productive, and the Trisomy 21 Model Atlas reinforces one of its central premises: triplicated genes generally behave in a predictable manner, with expression elevated close to the 1.5-fold expectation of a 3:2 copy number ratio in essentially every tissue, developmental stage, sex, and cell type in which they are expressed. Dosage sensitivity is, by this measure, a near-universal property of triplication rather than a feature of particular tissues or genes.

What the Atlas contributes is a systematic view of transcriptional dysregulation across the genome. Across seven tissues and three developmental stages in the Dp16 mouse model, and across five human iPSC-derived cell types, genes outside the triplicated region account for the overwhelming majority of differential expression, typically one to two orders of magnitude more genes than the triplication itself. These widespread and bidirectional downstream changes are largely distinct between tissues, with substantially different sets of DEGs in each context even where common pathways are affected. Understanding the transcriptional consequences of trisomy 21 therefore requires looking beyond the trisomy itself, to how each tissue and cell type responds to the elevated expression of a comparatively small set of triplicated genes, as well as the effects of accumulated structural, cellular, and functional changes that arise over the course of development. Mapping this complex landscape, alongside the dosage effect itself, is precisely what the Atlas is built to support.

The most striking feature of the dosage effect is how little it varies. Triplicated genes were elevated close to the 1.5-fold expectation in essentially every context examined, including across diverse tissues, developmental stages, sexes, and species, suggesting that the regulatory mechanisms shaping expression in each of these contexts do not, for the most part, offset the effect of copy number. The exceptions are correspondingly informative. Some triplicated genes are expressed in only a subset of tissues. A smaller number of triplicated genes showed fold-changes well above the dosage expectation, likely reflecting regulation by other dosage-sensitive factors or pathways. Amplification of this kind is more readily detected than suppression, since a triplicated gene expressed below its dosage expectation would typically remain elevated relative to wild-type.

Against this uniform backdrop, the scale of dysregulation beyond triplicated genes is the defining feature of the trisomic transcriptome. This imbalance has been evident in individual tissues and cell types, including our own earlier work^24,32^, but its consistency across such a broad panel of tissues, developmental stages, and species has not been directly assessed. These thousands of nontriplicated genes encompass most of the pathway-level signal and most of the tissue-specific variation observed downstream of the trisomy.

That downstream responses vary between tissues is expected. Mechanistic understanding depends on the composition of that variation: which genes change in which contexts, and which are shared. Interferon signaling, for example, was enriched in nearly every tissue and cell type examined, corroborating one of the most consistently reported features of Down syndrome across human cohorts and model systems^24,29,32^. However, the sets of genes contributing to that enrichment were distinct across contexts, and in some cases individual genes moved in opposite directions across tissues.

Pathway-level analyses are thus best treated as exploratory rather than definitive, particularly given that gene set annotations are themselves incomplete and evolving: shared enrichment can reflect related but distinct underlying transcriptional changes, an unavoidable property of summarizing a heterogeneous gene set in a single statistic. Resolving whether a shared annotation corresponds to a shared biological response requires inspecting the component genes across the contexts of interest, which the datasets in this release now make possible. The implication extends beyond mechanistic interpretation to the design of therapeutic interventions^30,32,41^: a strategy directed at a pathway enriched across multiple tissues may act on different genes, and to different effect, in each^14,24^.

Development, sex, and tissue pathology each modulate the downstream response while leaving the dosage signature intact. Cell-cycle-related genes illustrate the developmental dimension most clearly: strongly depleted at E18.5 but attenuated or reversed by later stages, consistent with a developmental dynamic rather than a persistent feature of the trisomic state, whereas elevated expression of interferon-related genes was evident at the earliest stage examined and persisted throughout. Sex contributes further variability in the downstream response, though reduced power at this sample size means genuine sex-dependent regulation cannot be distinguished from threshold effects. Sex remains an underexamined variable in Down syndrome research, and the patterns identified here indicate where larger, adequately powered analyses could be most informative.

Integrating histopathology adds a further dimension. Composite pathology scores reached significance in only a few tissues despite transcriptional dysregulation being detectable in essentially every tissue profiled. That disconnect may reflect the sensitivity of our composite scoring, the timeframe examined, or the considerable capacity of developing tissues to maintain functional architecture despite substantial molecular perturbation – a robustness essential for organismal viability. Longer-term cohorts and more sensitive structural assessments would help distinguish these possibilities.

A central question for any experimental model of Down syndrome is how well its findings generalize across model systems and to human biology. Data from iPSC-derived cell types differentiated from our panel of Human Trisome Project donor lines reproduced the same general organization observed in Dp16 tissues: largely consistent upregulation of triplicated genes across cell types, and a much larger set of nontriplicated DEGs differing substantially between them. Inflammatory gene sets, particularly IFNα and IFNγ responses, were enriched across nearly every cell type, again composed of related but distinct genes in each. That the same organization recurs in human cells, across five lineages spanning multiple germ layers and derived from twelve independent donors, indicates a general consequence of chromosome 21 triplication rather than a feature specific to the Dp16 model or to mouse biology. This expectation was reasonable but, until now, untested across this range of contexts.

The magnitude of the transcriptional response also varied markedly between iPSC-derived cell types, with only tens of DEGs detected in iAstrocytes and iNeurons compared with hundreds to thousands in the other cell types examined. Strongly cell-type-dependent differences in the number of DEGs have been reported previously in a microarray-based study of a panel of T21 and euploid neural progenitor cells, iPSCs, and source fibroblasts^27^, although the two studies differ markedly in the number of DEGs detected in undifferentiated iPSCs. Whether such differences reflect genuine variation in the sensitivity of particular cell types to T21, or technical factors including differentiation efficiency, culture heterogeneity, profiling platform, and statistical power, remains unclear.

### Limitations of the study

Several limitations should be considered when interpreting these results. Bulk transcriptomic profiling cannot distinguish cell-intrinsic changes in gene expression from shifts in the cellular composition of a tissue, and both are plausible contributors to the differences reported here – particularly in tissues where histopathological analysis identified altered tissue architecture. Resolving these contributions requires single-cell profiling. The sex-stratified and pathology-correlation analyses are the most power-limited in the Atlas: splitting each tissue by sex reduces group sizes to six animals, and the pathology correlations were performed within Dp16 animals only. Both should be treated as exploratory rather than definitive. In the case of sex, a within-sex resampling comparison would help determine whether the low overlap between male and female DEG sets reflects genuine sex-dependent regulation or the behavior of a significance threshold applied to small groups. Sequencing batch is partially confounded with tissue and timepoint for a subset of samples, though performing differential expression testing separately for each tissue-timepoint combination limits its influence on genotype comparisons.

This release is built on a single mouse model; Dp16 carries roughly two-thirds of the genes orthologous to HSA21 and no additional non-orthologous regions. Findings reported here may not extend to models with different genomic content. Developmental sampling is limited to three stages and histopathology to two, sufficient to identify broad developmental patterns but not to resolve trajectories in detail. The human arm relies on iPSC-derived cells, which are often considered developmentally immature relative to their primary counterparts and are cultured in isolation from tissue context, heterotypic cell-cell interactions, and circulating factors that shape gene expression *in vivo*. This is relevant to interpreting the cell-cycle results in particular, where the direction of pathway-level change varied markedly between cell types: whether this reflects differences in the proliferative state of these cultures or genuine context-dependence of the trisomy remains to be determined.

All datasets described here are available through the Experimental Models of Down Syndrome portal within the INCLUDE Data Hub (https://experimentalmodels.includedcc.org/trisomy-21-model-atlas.html), where they can be queried and visualized by organ, cell type, or gene. A researcher working on a candidate gene can see where it is expressed across seven mouse tissues and five human cell types, whether it responds to trisomy in each, and how that response varies with developmental stage and sex – without generating new data or reanalyzing raw sequencing files. Complete differential expression results are available for reanalysis under different thresholds or analytical frameworks, and raw sequencing data are deposited in GEO under SuperSeries GSE347972.

This release is intended as a foundation rather than a completed resource. Work in progress includes single-cell transcriptomics, which will address the cell-composition ambiguity inherent to bulk profiling, and multiplexed imaging to extend histopathological characterization beyond composite scoring. Additional mouse models and iPSC-derived systems will broaden the range of genomic content and cellular contexts represented. Data will be released through the Experimental Models portal as they are generated, under the metadata standards and persistent identifiers established here.

More broadly, the Experimental Models portal (https://experimentalmodels.includedcc.org) is designed to host datasets from across the Down syndrome research community, and the Atlas represents one contribution to that shared infrastructure. Datasets from other groups that extend or complement this work – different models, tissues, timepoints, or modalities – can be accommodated within the same framework, and surplus biospecimens from the cohorts described here are available on request through the portal (https://experimentalmodels.includedcc.org/virtualbiorepository.html).

Trisomy of chromosome 21 has been recognized as the cause of Down syndrome since 1959^42^, and its approximate gene content has been known for over two decades^43^. What has proven harder to establish is what happens next: which genes are dysregulated in which tissues, how that dysregulation accumulates through development and across the lifespan, and how closely the resulting landscape is recapitulated across experimental model systems. The Trisomy 21 Model Atlas aims to help answer these questions systematically. Against the consistent backdrop of dosage-driven upregulation, the substantial majority of transcriptional change occurs outside the triplicated region and varies markedly with biological context. Characterizing that context-dependence is a prerequisite for connecting chromosome 21 gene dosage to the phenotypes and co-occurring conditions experienced by people with Down syndrome^44^ – a central aim of the INCLUDE Project^45^ – and for the design of effective and personalized therapeutic interventions aimed at improving health and lifespan in this population^19^.

## RESOURCE AVAILABILITY

### Lead contact

Further information and requests for resources and reagents should be directed to and will be fulfilled by the lead contact, Matthew D. Galbraith.

### Materials availability

Human iPSC lines generated through the Crnic Institute Human Trisome Project and used in this study are available from the lead contact upon request, subject to a completed Materials Transfer Agreement and the terms of the originating study protocol (COMIRB 15-2170). The Dp(16)1Yey mouse strain is available from The Jackson Laboratory (JAX:013530). Surplus biospecimens from the mouse cohorts described here are available on request through the Experimental Models of Down Syndrome portal (https://experimentalmodels.includedcc.org/virtualbiorepository.html). This study did not generate new unique reagents.

### Data and code availability

Bulk RNA-sequencing data have been deposited at GEO and are publicly available. Accession numbers are listed in the key resources table. Data generated for this study are available under SuperSeries accession GSE347972, comprising SubSeries for each mouse tissue–timepoint combination and for undifferentiated iPSCs, iHSPCs, and iMonocytes. iHepatocyte data were previously reported in Dunn et al.^14^ (GSE296449), and iNeuron and iAstrocyte data are reported in Dooling et al.^36^ (GSE344531 and GSE344533). Per-series accessions and sequencing metrics for all datasets are provided in **Table S1**. Processed data and differential expression results are additionally available through the Experimental Models of Down Syndrome portal within the INCLUDE Data Hub (https://experimentalmodels.includedcc.org/trisomy-21-model-atlas.html).

All original code has been deposited on GitHub (https://github.com/Linda-Crnic-Institute-for-Down-Syndrome/t21-model-atlas-analysis) and archived at Zenodo (https://doi.org/10.5281/zenodo.23198762) and is publicly available.

Any additional information required to reanalyze the data reported in this paper is available from the lead contact upon request.

## ACKNOWLEDGEMENTS

This work was supported primarily by NIH grant R24OD035579 (M.D.G., K.D.S., J.M.E.) as part of the INCLUDE Project at the Office of the Director. Additional support was provided through NIH grant U2CHL156291 (M.D.G. and J.M.E) and U24AG092191 (M.D.G. and J.M.E), the Linda Crnic Institute for Down Syndrome, the Global Down Syndrome Foundation, the Anna and John J. Sie Foundation, and the Boettcher Foundation (K.D.S.).

## AUTHOR CONTRIBUTIONS

K.R., data curation, formal analysis, investigation, visualization, methodology, software, writing – original draft, writing – review and editing; A.T., investigation, visualization, methodology, writing – original draft, writing – review and editing; B.N., investigation, visualization, methodology, writing – original draft, writing – review and editing; K.S., investigation; L.D., investigation; M.C.H., investigation; R.A.H., investigation; S.C., investigation; A.L.R., resources; J.M.E., supervision, funding acquisition, and writing – review and editing; K.D.S., conceptualization, supervision, funding acquisition, visualization, methodology, and writing – review and editing; M.D.G., conceptualization, supervision, funding acquisition, visualization, methodology, software, writing – original draft, and writing – review and editing

## DECLARATION OF INTERESTS

J.M.E. has provided consulting services to Eli Lilly and Co., Gilead Sciences Inc., Biohaven Pharmaceuticals, Perha Pharmaceuticals, and AbbVie. K.D.S. has provided consulting services to AbbVie. All other authors declare that they have no competing interests.

## DECLARATION OF GENERATIVE AI AND AI-ASSISTED TECHNOLOGIES IN THE MANUSCRIPT PREPARATION PROCESS

During the preparation of this work, the authors used AI-assisted technologies to improve grammar and clarity.

## SUPPLEMENTAL INFORMATION TITLES AND LEGENDS

1. Document S1. Figure S1-S7
2. Table S1. Experimental design, sample numbers, and RNA-seq QC for mouse and iPSC experiments.
3. Table S2. Dp16 versus WT differential expression analysis results across all tissues from the E18.5, 1-month, and 4-month group.
4. Table S3. Dp16 vs WT GSEA results from all tissues in 4mo group.
5. Table S4. Dp16 vs WT GSEA results from all tissues in E18.5 and 1-month group.
6. Table S5. 4-month vs 1-month differential expression analysis results from Dp16 and WT lungs.
7. Table S6. Dp16 vs WT differential expression analysis results from all tissues in females from 4mo group and from all tissues in males from 4mo group
8. Table S7. Dp16 vs WT GSEA results from all tissues in females from 4mo group and from all tissues in males from 4mo group
9. Table S8. Histopathology scores from E18.5 mice.
10. Table S9. Histopathology scores from 1mo mice.
11. Table S10. Histopathology scores from 4mo mice.
12. Table S11. Histopathology scores versus bulk RNAseq from 4mo kidney and lung.
13. Table S12. T21 vs D21 differential expression analysis results from iPSCs.
14. Table S13. T21 vs D21 GSEA results from iPSCs.
15. Table S14. Scoring rubric for 1mo and 4mo histopathology

## STAR METHODS

### Key Resources Table

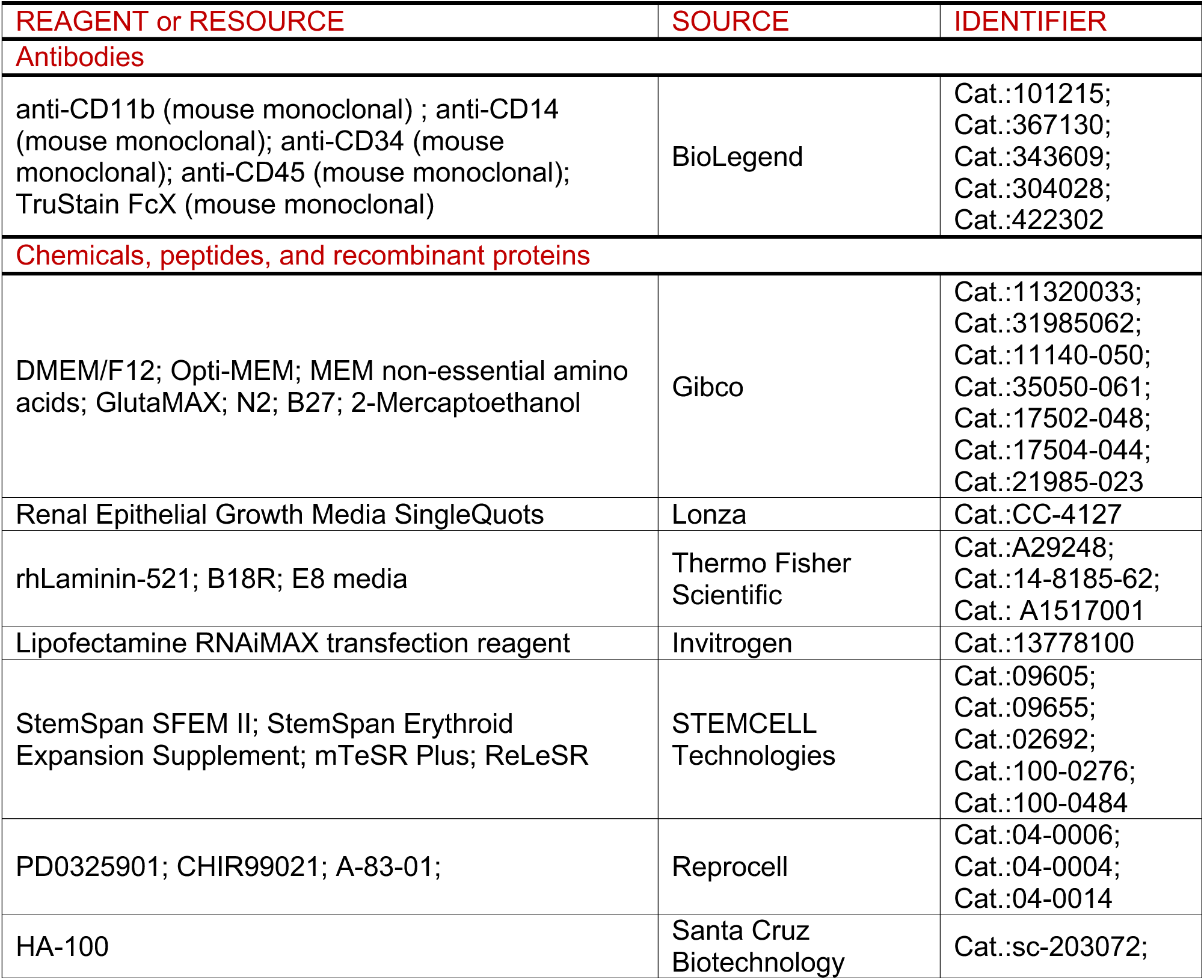
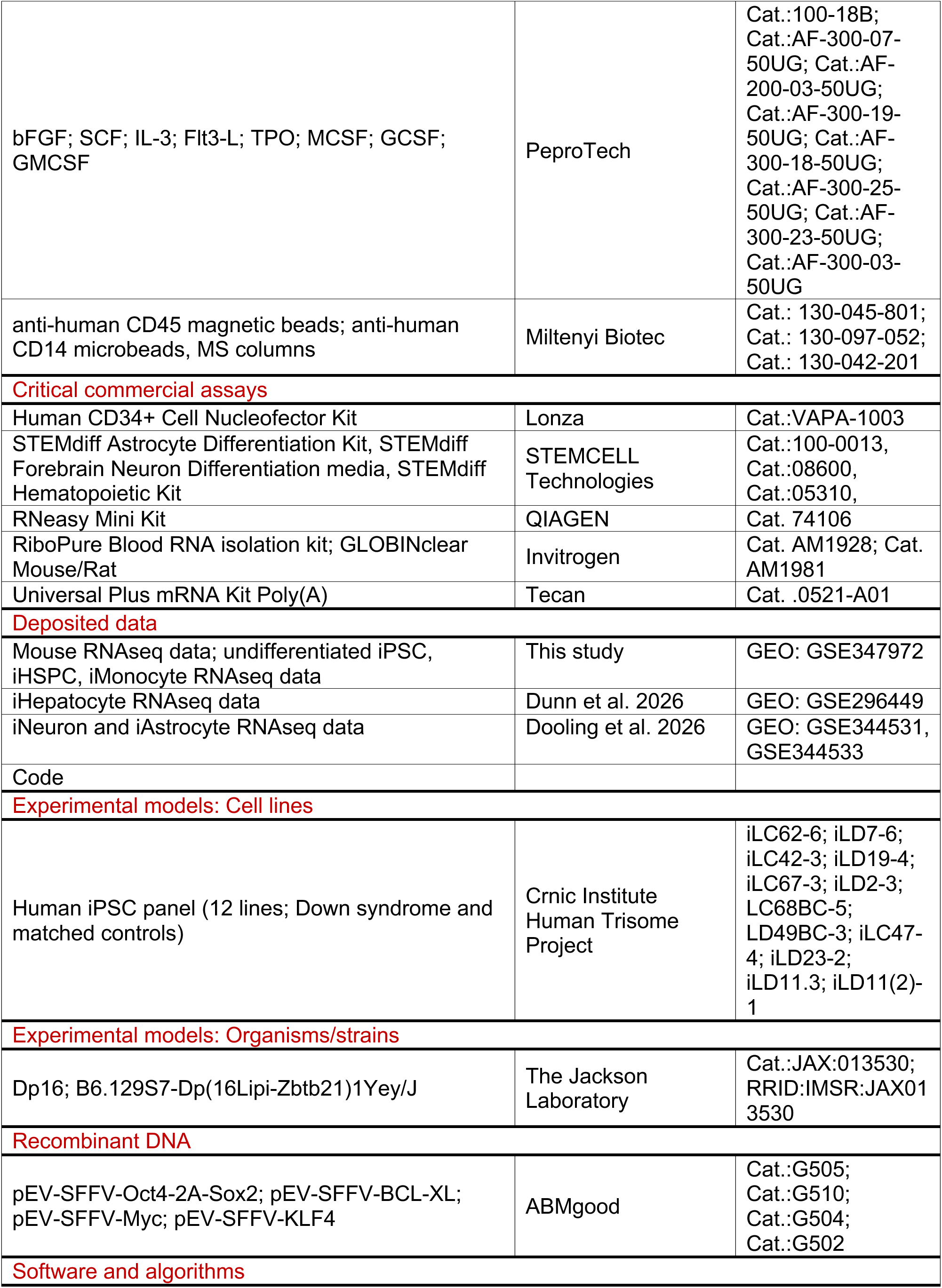
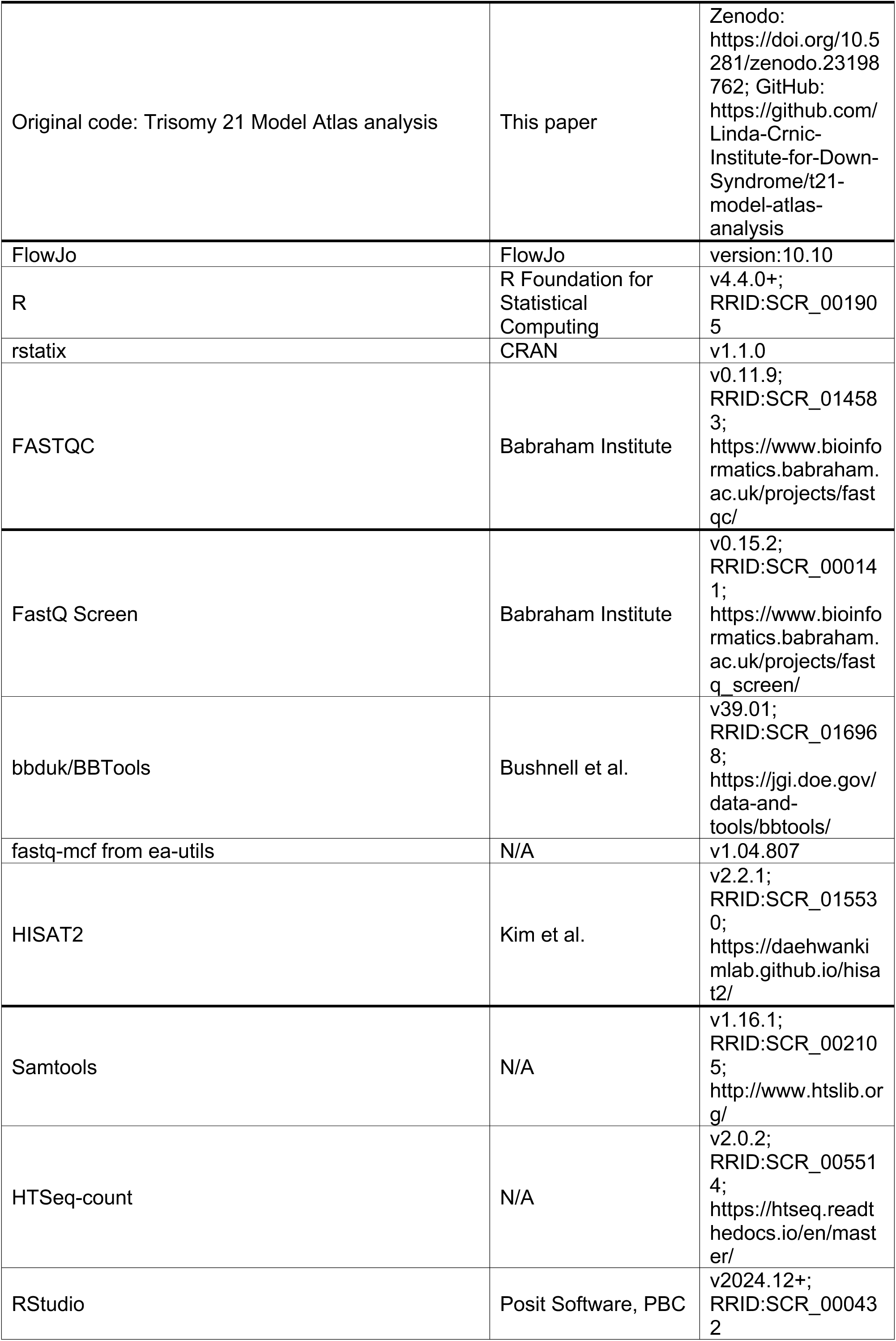
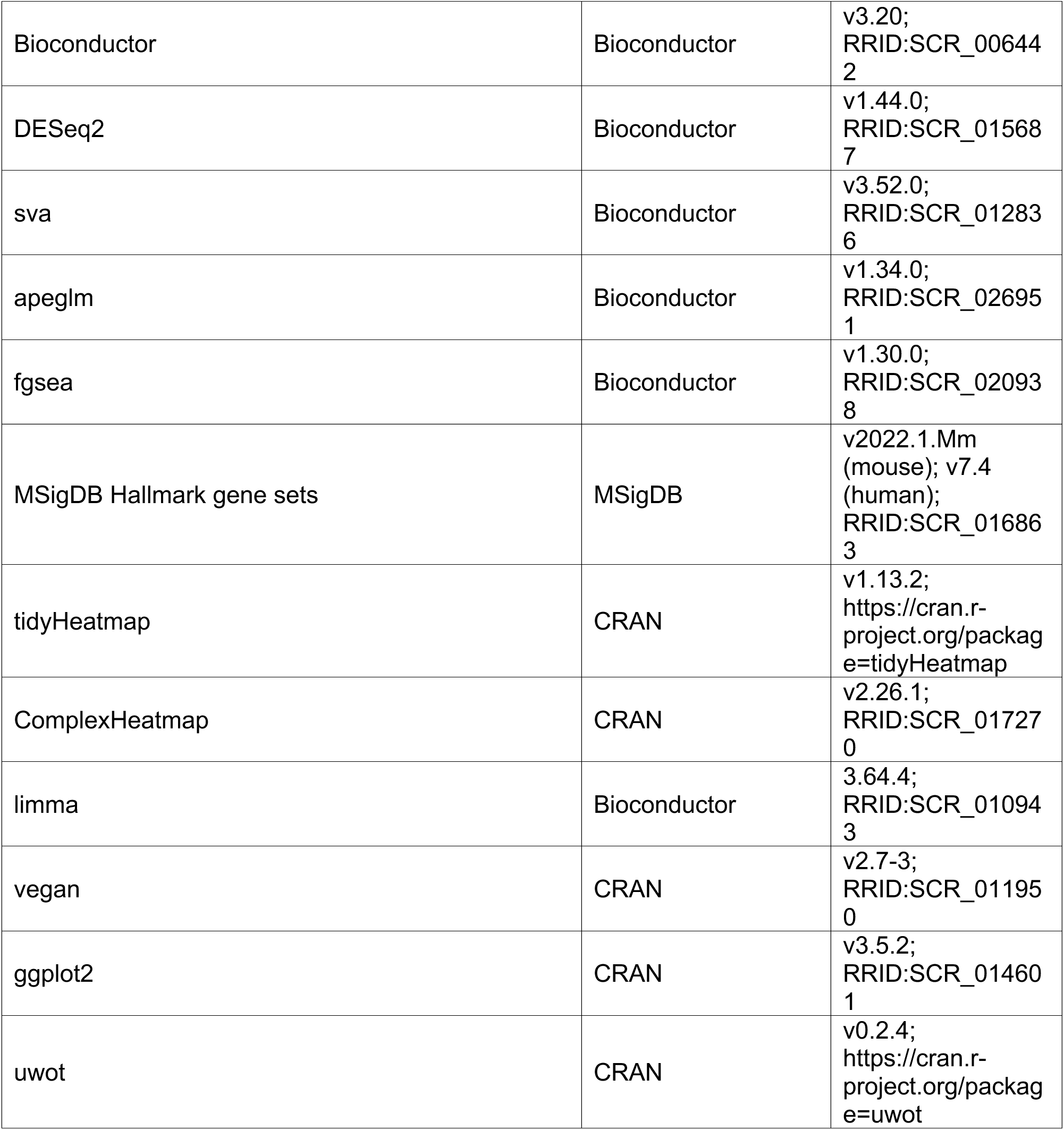

## Experimental model and study participant details

### Experimental design

Mouse cohorts were balanced by genotype and sex at each timepoint, with 12 WT and 12 Dp16 mice per timepoint (6 female and 6 male per genotype). For the 1- and 4-month cohorts, 12 tissue types were collected from each mouse for histopathological evaluation, and seven matched biospecimens from the same animals were collected for RNA-seq (whole blood, brain, lung, heart, liver, kidney, and large intestine). Thus, within each postnatal timepoint, histopathology and transcriptomic datasets were matched at the animal level. This design minimized variability attributable to differences in the animals contributing to each dataset and preserved matched sampling across tissues. For the E18.5 cohort, whole embryos were collected for histopathological evaluation, whereas brain, heart, lung, liver, whole intestine, and kidney were collected from fresh embryos for RNA-seq. For analyses across developmental stages, E18.5 whole intestine was grouped with large intestine from the 1- and 4-month cohorts. Individual mice were considered as biological replicates. Samples that did not meet sequencing quality control criteria were excluded from downstream analyses; final sample numbers for each tissue-timepoint are provided in **Table S1**.

The human iPSC study comprised 12 independent donor-derived iPSC lines, including 6 D21 and 6 T21 lines, with 2 female and 4 male donors per genotype. Individual iPSC lines were considered as biological replicates. The same panel of lines was used across the iPSC-derived cell types with final sample numbers for each cell type provided in **Table S1**.

### Animal husbandry

Experiments were approved by the Institutional Animal Care and Use Committee at the University of Colorado Anschutz Medical Campus under protocol #00111 and performed in accordance with National Institutes of Health (NIH) guidelines. Dp16 (B6.129S7-Dp(16Lipi-Zbtb21)1Yey/J) mice were originally purchased from the Jackson Laboratory and maintained on the C57BL/6J background in specific pathogen–free conditions. Mice were housed separately by sex in groups of one to five mice per cage under a 14-hour light:10-hour dark cycle with controlled temperature and 35% humidity and had ad libitum access to food (16 kcal% fat diet) and water. For genotyping, genomic DNA was prepared from 1 to 2 mm of tail or ear tissue for automated genotyping by quantitative PCR with specific probes designed against *Lipi* and *Zbtb21* (Transnetyx, Cordova, TN, USA). Upon termination of the studies, all animals were euthanized by CO_2_ asphyxiation and cardiac puncture, then immediately perfused with 1x PBS.

### Mouse postnatal tissue collection

Upon euthanasia, 1-month and 4-month mouse tissues were collected for transcriptome profiling (RNA-seq) and histopathological analysis. Whole blood was collected via cardiac puncture in EDTA and 500 µL of whole blood was mixed with 1.3 mL RNAlater (Invitrogen, Carlsbad, CA, USA) and stored at −80°C until RNA extraction. Whole blood is referred to as blood throughout. Organs were distributed for RNA-seq or histopathology as follows: the brain and lung were cut mid-sagittally with the right half going for RNA-seq and left half for histopathology. The heart was cut transversely with the top going for RNA-seq and the bottom for histopathology. For the liver the medial lobe was used for histopathology and the left lobe was used for RNA-seq. The right kidney was used for RNA-seq and the left for histopathological analysis. The large intestine was flushed with 1x PBS and the proximal and distal ends were cut for RNA-seq and the entire middle section was fixed for histopathological analysis. Adipose tissue was collected from the abdomen, quadriceps muscle was collected as representative skeletal muscle, and shaved dorsal skin and spleens were collected for histopathological analysis. The small intestine was flushed with 1× PBS before fixation for histopathological analysis. Finally, femurs were dissected, cleaned of surrounding soft tissue, and fixed for histopathological evaluation, including bone marrow and bone grading.

### Mouse embryo tissue collection

Male Dp16 mice were crossed overnight with 8-to 12-week-old estrus-synchronized female C57BL/6J mice. Dams were checked daily for vaginal plugs; the morning of first visual confirmation was denoted embryonic day (E) 0.5. At E18.5, embryos were collected after CO_2_ asphyxiation and cervical dislocation of dams and allowed to exsanguinate on ice in 1x PBS. For histopathology, tails were collected for genotyping (Transnetyx) and whole embryos were fixed in 10% neutral buffered formalin for 48 hours before being cut mid-sagittally and both halves embedded in paraffin. For RNA-seq profiling, organs including brain, heart, lung, liver, whole intestine, and kidney from E18.5 were manually dissected from fresh embryos and then flash-frozen and stored at −80°C. Whole intestine are grouped with large intestine from 1-month and 4-month age groups throughout.

### Human iPSC generation and maintenance

A panel of 12 iPSC lines were used for this study, comprising 6 D21 and 6 T21 lines (see **Table S1**). These were previously generated from age-and sex-matched individuals with and without Down syndrome from Crnic Institute’s Human Trisome Project (HTP) under a study protocol approved by the Colorado Multiple Institutional Review Board (COMIRB 15–2170) and ten of these twelve lines have been described in previous publications^14,46–48^. The iPSCs were generated from either urine-derived renal epithelial cells (RECs) or peripheral blood mononuclear cells (PBMCs) through the Gates Center for Regenerative Medicine. RECs were reprogrammed into iPSCs using a cocktail of miRNAs-367/302s and mRNAs encoding six reprogramming factors: a modified version of *Oct4* fused with the *MyoD* transactivation domain (called M3O), *Sox2*, *Klf4*, *cMyc*, *Lin28A*, and *Nanog*^49^. Briefly, RECs were expanded in renal epithelial growth media (REGM) consisting of DMEM/F12 media (Gibco, Grand Island, NY, USA, Cat. 11320033) supplemented with Renal Epithelial Growth Media SingleQuots (Lonza, Basel, Switzerland, Cat. CC-4127). After expansion, RECs were plated in 6-well plates coated with rhLaminin-521 (ThermoFisher, Waltham, MA, USA, Cat. A29248) at 1e5 cells per well. RECs were transfected with miRNAs and mRNAs on days 1, 3, 5, 7, 9, 11, and 13 post-plating using Opti-MEM (Gibco, Cat. 31985062) supplemented with Lipofectamine RNAiMax transfection reagent (Invitrogen, Cat. 13778100). After transfection, RECs were cultured in REGM supplemented with B18R (ThermoFisher, Cat. 14-8185-62). From day 7 onward, REGM was supplemented with bFGF (Peprotech, Cranbury, NJ, USA, Cat. 100-18B). On Day 9, REGM was replaced with E8 media (ThermoFisher, Cat. A1517001).

PBMCs were reprogrammed to generate iPSCs using an episomal approach. Briefly, cryopreserved PBMCs were thawed, and erythroid progenitors were allowed to expand for 7 days in StemSpan SFEM II Media (StemCell Technologies, Vancouver, Canada, Cat. 09605) supplemented with StemSpan Erythoid Expansion Supplement (StemCell Technologies, Cat. 02692). On day 7, cells were nucleofected using the Human CD34+ Cell Nucleofector Kit (Lonza, Cat. VAPA-1003) and the following vectors: pEV-SFFV-Oct4-2A-Sox2 (ABMgood, Vancouver, Canada, Cat. G505), pEV-SFFV-BCL-XL (ABMgood, Cat. G510), pEV-SFFV-Myc (ABMgood, Cat. G504), and pEV-SFFV-KLF4 (ABMgood, Cat. G502). Cells were then cultured in reprogramming media consisting of StemSpan SFEM II media supplemented with MEM non-essential amino acids (Gibco, Cat. 11140-050), GlutaMAX (Gibco, Cat. 35050-061), N2 (Gibco, Cat. 17502-048), B27 (Gibco, Cat. 17504-044), 2-Mercaptoethanol (Gibco, Cat. 21985-023), PD0325901 (Reprocell, Beltsville, MD, USA, Cat. 04-0006), CHIR99021 (Reprocell, Cat. 04-0004), A-83-01 (Reprocell, Cat. 04-0014), HA-100 (Santa Cruz Biotechnology, Dallas, TX, USA, Cat. sc-203072), and bFGF. On days 12-15 post-nucleofection, media was gradually transitioned to E8 media starting at 25% E8 media on day 12 and increasing an additional 25% daily with each media change.

Once reprogrammed, the iPSCs were maintained on Matrigel-coated plates (Corning Incorporated, Corning, NY, USA Cat. 354277) in mTeSR Plus media (StemCell Technologies, Cat. 100–0276). Spontaneously differentiating cells were marked and manually removed. iPSCs were routinely passaged as aggregates using ReLeSR (StemCell Technologies, Cat. 100-0484). Mycoplasma testing performed through the Functional Genomics Shared Resource at the University of Colorado indicated no mycoplasma contamination. Genomic integrity and retention of the triplicated chromosome 21 in the iPSCs were validated by g-banded karyotype analysis through the University of Colorado Cancer Center Pathology Shared Resource - Cytogenetics Section.

### iPSC differentiation

iPSCs were differentiated into iHepatocytes over the course of 15 days as previously reported^14,50^. Differentiation was validated in the original publication by immunofluorescence staining of albumin. iAstrocytes and iNeurons were differentiated as previously described (Dooling et al., 2026) using the STEMdiff™ Astrocyte differentiation Kit (STEMCELL technologies, Cat. 100-0013) and STEMdiff™ Forebrain Neuron Differentiation media (STEMCELL technologies, Cat. 08600). iAstrocytes and iNeuron differentiation was validated by detection of GFAP (iAstrocytes) and MAP2 and NeuN (iNeurons) using immunofluorescence^36^.

iPSCs were differentiated into iHematopoietic Stem and Progenitor Cells (iHSPCs) using the STEMdiff™ Hematopoietic Kit (STEMCELL technologies, Cat. 05310) according to the manufacturer’s protocol. Cells were plated in aggregates in 6-well plates with ∼130 aggregates per well. Cells were cultured in Media A from days 0-3, with a half media change on day 2. On day 3 media was fully changed to Media B and cells were cultured until day 12 with half media changes on days 5, 7, and 10. On day 12, non-adherent cells were collected. Prior to RNA isolation for RNA-sequencing, hematopoietic cells were enriched using anti-human CD45 magnetic beads (Miltenyi Biotec, Gaithersburg, MD, USA, Cat. 130-045-801) and MS columns (Miltenyi Biotec, Cat. 130-042-201) according to the manufacturer’s instructions.

iPSC-derived monocytes (iMonocytes) were differentiated from iHSPCs described above. After day 12 of iHSPC differentiation, wells were washed with DMEM/F12 to detach any non-adherent cells. These non-adherent iHSPCs were resuspended in myeloid differentiation media consisting of StemSpan SFEM II (STEMCELL technologies, Cat. 09655) supplemented with 50 ng/mL SCF (PeproTech, Cat. AF-300-07-50UG), 50 ng/mL IL-3 (PeproTech, Cat. AF-200-03-50UG), 50 ng/mL Flt3-L (PeproTech, Cat. AF-300-19-50UG), 4 ng/mL TPO (PeproTech, Cat. AF-300-18-50UG), 50 ng/mL MCSF (PeproTech, Cat. AF-300-25-50UG), 50 ng/mL GCSF (PeproTech, Cat. AF-300-23-50UG), and 25 ng/mL GMCSF (PeproTech, Cat. AF-300-03-50UG). Cells were resuspended at 5e5 cells/mL in myeloid differentiation media and plated in 6-well plates with up to 4 mL per well. Cells were cultured for 6 days with half media changes performed every 48 hours. Monocytes were enriched using anti-human CD14 microbeads (Miltenyi Biotec, Cat. 130-097-052) and MS columns (Miltenyi Biotec, Cat. 130-042-201) according to the manufacturer’s instructions.

### Method Details

#### Histopathology of mouse tissues

All histopathological mouse tissues were fixed in 10% formalin for 24 hours prior to processing and paraffin embedding. Femurs were fixed for 24 hours and then decalcified with EDTA prior to processing and embedding for bone marrow histopathology. Formalin-fixed paraffin-embedded (FFPE) tissues were sectioned at 5 microns and stained with hematoxylin and eosin (H&E) and femurs were also stained with Giemsa and periodic acid–Schiff (PAS). Blinded veterinary anatomic pathologists assessed and graded all tissues according to previously established histopathological grading schemes unique to each tissue.

Individual pathological features were assigned scores reflecting the severity and extent of pathology, with higher scores indicating greater abnormalities (see **Table S14**). For each tissue, component scores were summed to generate a composite score representing the overall severity of tissue pathology. Histopathology scores from Dp16 and wild-type (WT) mice were analyzed at E18.5, 1 month, and 4 month separately.

Composite scores were compared between Dp16 and wild-type (WT) mice at E18.5, 1 month, and 4 months of age using unpaired Wilcoxon rank-sum tests. For 1- and 4-month-old mice, tests were performed separately for each tissue using a one-sided alternative hypothesis that Dp16 mice would exhibit greater pathology than WT mice. Comparisons with fewer than two observations per genotype or with no variation in scores were excluded. Postnatal p-values were not adjusted for multiple testing. All analyses were performed in R using the rstatix package (v1.1.0).

#### Flow cytometry

Flow cytometry was performed on days 12 and 15 of differentiation for iHSPCs and iMonocytes, respectively. Cells were collected and washed three times in flow buffer (97.5% dPBS and 2.5% BSA). Cells were labeled with antibodies against CD11b (Biolegend, San Diego, CA, USA, Cat. 101225), CD14 (Biolegend, Cat. 367130), CD34 (Biolegend, Cat. 343609), CD45 (Biolegend, Cat. 304028), and Fc block (Biolegend, Cat. 422302) in 100 µl of flow buffer at 4°C for 20 minutes. Samples were washed three times in flow buffer then resuspended in flow buffer with 1 µg/ml propidium iodide (ThermoFisher, Cat. P3566) before data acquisition on a five-laser Cytek Aurora spectral flow cytometer. Flow cytometry analysis was performed using FlowJo software v10.10.

#### Bulk RNA-sequencing

Upon euthanasia, 1-month and 4-month mouse tissues were immediately collected and placed in 594 μL of lysis buffer RLT Plus (QIAGEN, Redwood City, CA, USA) and 6 μL of 2-mercaptoethanol in Lysing Matrix D tubes (MP Biomedicals, Santa Ana, CA, USA), homogenized, then stored at −80°C. Embryonic tissues were flash frozen at time of harvest. Prior to extraction embryonic tissues were homogenized in 346.5 μL RLT Plus and 3.5 μL 2-mercaptoethanol. Whole blood was collected via cardiac puncture at time of euthanasia and collected into EDTA K3 tubes (Sarstedt, Nümbrecht, Germany), then transferred to 1.3 mL RNAlater and frozen.

For mouse tissues and human iPSC cell lines, total RNA was isolated using the RNeasy Mini Kit (QIAGEN, Cat. 74106) following the manufacturer’s instructions. Whole blood total RNA was extracted using the mouse RiboPure blood RNA isolation kit (Invitrogen, Cat. AM1928) followed by Globin mRNA removal using the GLOBINclear Mouse/Rat (Invitrogen, Cat. AM1981) kit following manufacturer guidelines. RNA quality was assessed using an Agilent 2200 TapeStation. Strand-specific libraries were prepared using a Universal Plus mRNA Kit Poly(A) (Tecan, Männedorf Switzerland, Cat. 0521-A01). Libraries were sequenced on an Illumina NovaSeq 6000 (150-bp, paired-end) at the CU Anschutz Genomics and Microarray Core, yielding approximately 28M-86M sequences per library (see **Table S1**).

Data quality was assessed on a per-sample basis using FASTQC (v0.11.9) (https://www.bioinformatics.babraham.ac.uk/projects/fastqc/) and FastQ Screen (v0.15.2, https://www.bioinformatics.babraham.ac.uk/projects/fastq_screen/). Low-quality reads were filtered and trimmed with bbduk from BBTools (v39.01) and fastq-mcf from ea-utils (v1.04.807, https://expressionanalysis.github.io/ea-utils/). Mouse transcriptomes were aligned to the mouse reference genome (GRCm38) using HISAT2 (v2.2.1) in paired, spliced-alignment mode with a GRCm38 index with Gencode M24 annotation GTF. iPSC transcriptomes were aligned to the human reference genome (GRCh38) using HISAT2 (v2.2.1) in paired, spliced-alignment mode with a GRCh38 index with Gencode v33 annotation GTF. Alignments were sorted and filtered for mapping quality (MAPQ>10) using Samtools (v1.16.1). Gene-level count data were quantified using HTSeq-count (2.0.2) with the following options (--stranded=reverse -minaqual=10 -type=exon --mode=intersection-nonempty).

### Quantification and statistical analysis

Statistical analyses and plot generation for all datasets was performed using R (R 4.4.0+/RStudio 2024.12+/Bioconductor 3.20)^51,52^. All p-values calculated from RNA-sequencing comparisons were adjusted using the Benjamini–Hochberg method, with statistical significance defined as q < 0.10.

For RNA-seq data preprocessing and all differential expression analyses, genes were evaluated if they had >0.5 counts per million (CPM) in at least half of the samples being evaluated. Differential gene expression was evaluated using DESeq2 v1.44.0^53^ with surrogate variables determined by svaseq() from the sva R package 3.52.0 as covariates (SVA)^54^. Surrogate variables were estimated from DESeq2-normalized counts while protecting the variable of interest, with the number of surrogate variables determined automatically. Log_2_ fold-change estimates were shrunk using apeglm; these shrunken estimates are designated as log_2_FC_adj, with FC_adj representing the corresponding fold change on the linear scale (2^log_2_FC_adj). Unshrunken DESeq2 estimates are designated as log_2_FC and FC, respectively. P-values were calculated from the unshrunken DESeq2 estimates and padj represents q-values. Differential expression analyses were performed separately for each tissue-timepoint, and separately for each sex-tissue-timepoint, comparing Dp16 with WT samples within each dataset. Differential expression analyses comparing timepoints were performed separately for WT and Dp16 samples. Associations between gene expression and histopathology were evaluated separately within each tissue-timepoint among Dp16 mice by modeling composite histopathology scores as a continuous predictor, with SVA-derived surrogate variables as covariates. The corresponding log_2_ fold change represents the estimated expression change per unit increase in histopathology score. For comparisons evaluating whether triplicated-gene expression in Dp16 mice differed from WT by more than 1.5-fold, the DESeq2 lfcThreshold argument was set to log_2_(1.5), using the default two-sided altHypothesis = “greaterAbs” test to determine whether the absolute log_2_ fold change exceeded this threshold.

Gene set enrichment analysis (GSEA)^55^ was performed with the fgsea package v1.30.0^56^ using species-specific Molecular Signatures Database (MSigDB) Hallmark gene set collections (v2022.1.Mm for mouse datasets and v7.4 for human datasets). Genes were ranked, by apeglm-shrunken log_2_ fold changes from the corresponding SVA-adjusted DESeq2 comparison and enrichment was assessed using an unweighted statistic. Hallmark gene sets with a Benjamini-Hochberg-adjusted q < 0.1 were considered significant.

Heatmaps were generated using the tidyHeatmap (v1.13.2) and ComplexHeatmap (2.26.1) R packages. Volcano plots were generated using a custom R function. For specific visualizations, transcripts per million (TPM) values were calculated from gene-level counts by normalizing counts to summed non-overlapping exonic gene lengths derived from the GTF annotation and scaling the resulting values within each sample to a total of one million. TPM values were log_2_-transformed after addition of a pseudocount (log_2_(TPM + 1)), and the estimated effects of surrogate variables were removed using limma::removeBatchEffect. The adjusted expression values were standardized to z-scores for specific genes across samples. Mean z-scores were then calculated within each tissue and genotype and used to determine bubble size in the visualization. Normalized counts were calculated by DESeq2. Pairwise overlap of differentially expressed gene sets across tissues was quantified using the Jaccard similarity index and calculated separately for triplicated and nontriplicated genes using the vegan R package (v2.7-3). Pairwise Jaccard similarity between gene sets was defined as the size of their intersection divided by the size of their union. Pairwise Spearman rank correlations of adjusted log_2_(fold-change) from differentially expressed genes were computed separately for triplicated and nontriplicated genes. GSEA enrichment plots were generated using a custom R function. Sina plots showing all points jittered horizontally by local density with bars representing medians were generated using the ggplot2 (3.5.2) R package. For symmetric visualization of gene expression in female mice versus male mice within a genotype-tissue-timepoint group, fold-change ratios that were less than one were transformed as -1/fold change.

Uniform Manifold Approximation and Projection (UMAP) dimensionality reduction was performed on variance-stabilizing transformed (VST) counts from the full set of mouse bulk RNA-seq samples, without covariate adjustment. Principal component analysis was performed first, either on all expressed genes passing the low-expression filter or restricted to triplicated genes, and the first 30 principal components were used as input to UMAP. Embeddings were computed with the uwot R package using n_neighbors = 15, min_dist = 2, Euclidean distance, and a fixed random seed (123).

## Notes

https://experimentalmodels.includedcc.org/trisomy-21-model-atlas.html

### Summary of Updates

Minor updates to Figures 2B and 7A,B. Methods reorganized into a key resources table and separate experimental, method, and statistical sections; descriptions of the UMAP analysis and dosage threshold test added; author contributions, keywords, and a resource availability section added; references expanded from 47 to 56 entries. Analysis code deposited on GitHub and archived at Zenodo. Data, analyses, and conclusions are unchanged.

