## Supplemental Document S1 for "A Trisomy 21 Model Atlas reveals conserved dosage effects and context-specific transcriptional responses across mouse and human models of Down syndrome"

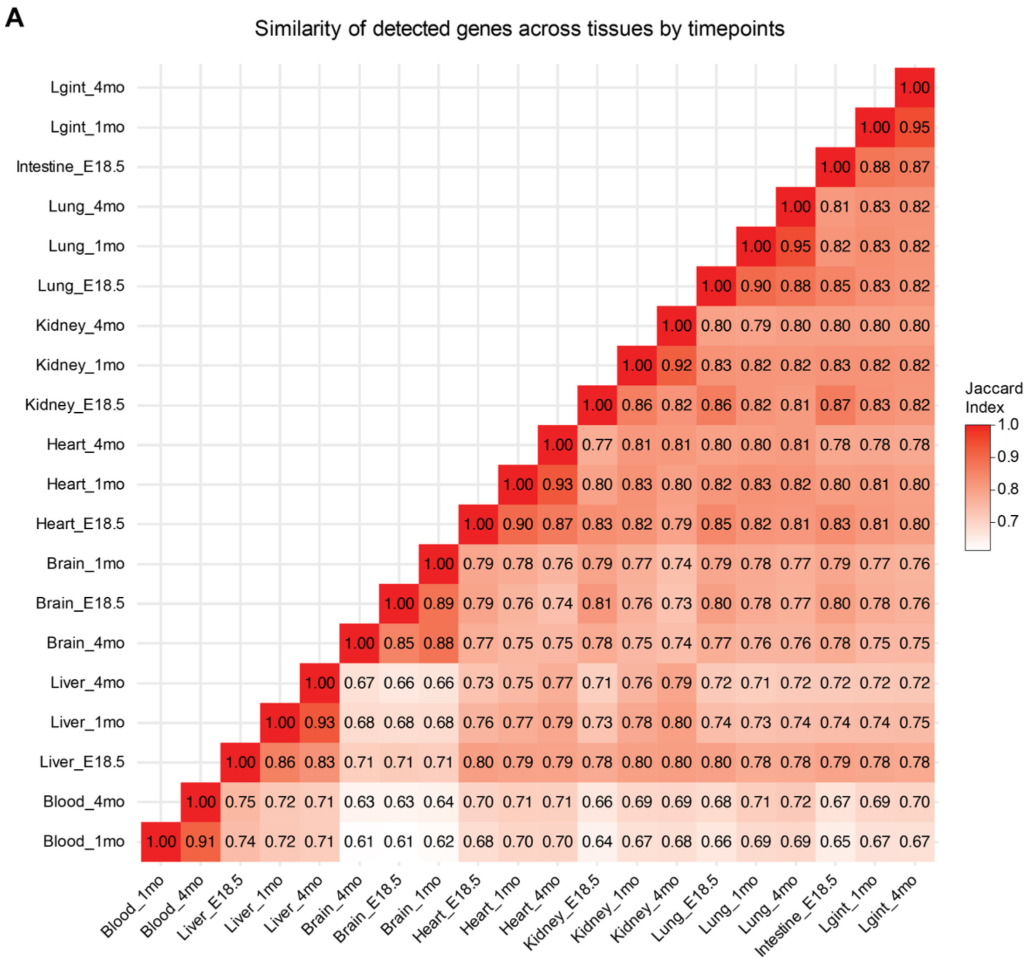


**Supplementary Figure 1, related to Figure 1. Overview of the Trisomy 21 Model Atlas study design.** Pairwise Jaccard index of detected genes between tissue–timepoint combinations across all seven mouse tissues and three developmental timepoints (12 WT and 12 Dp16 mice per tissue-timepoint). Intestine was collected as whole intestine at E18.5 and as large intestine at 1 and 4 months.

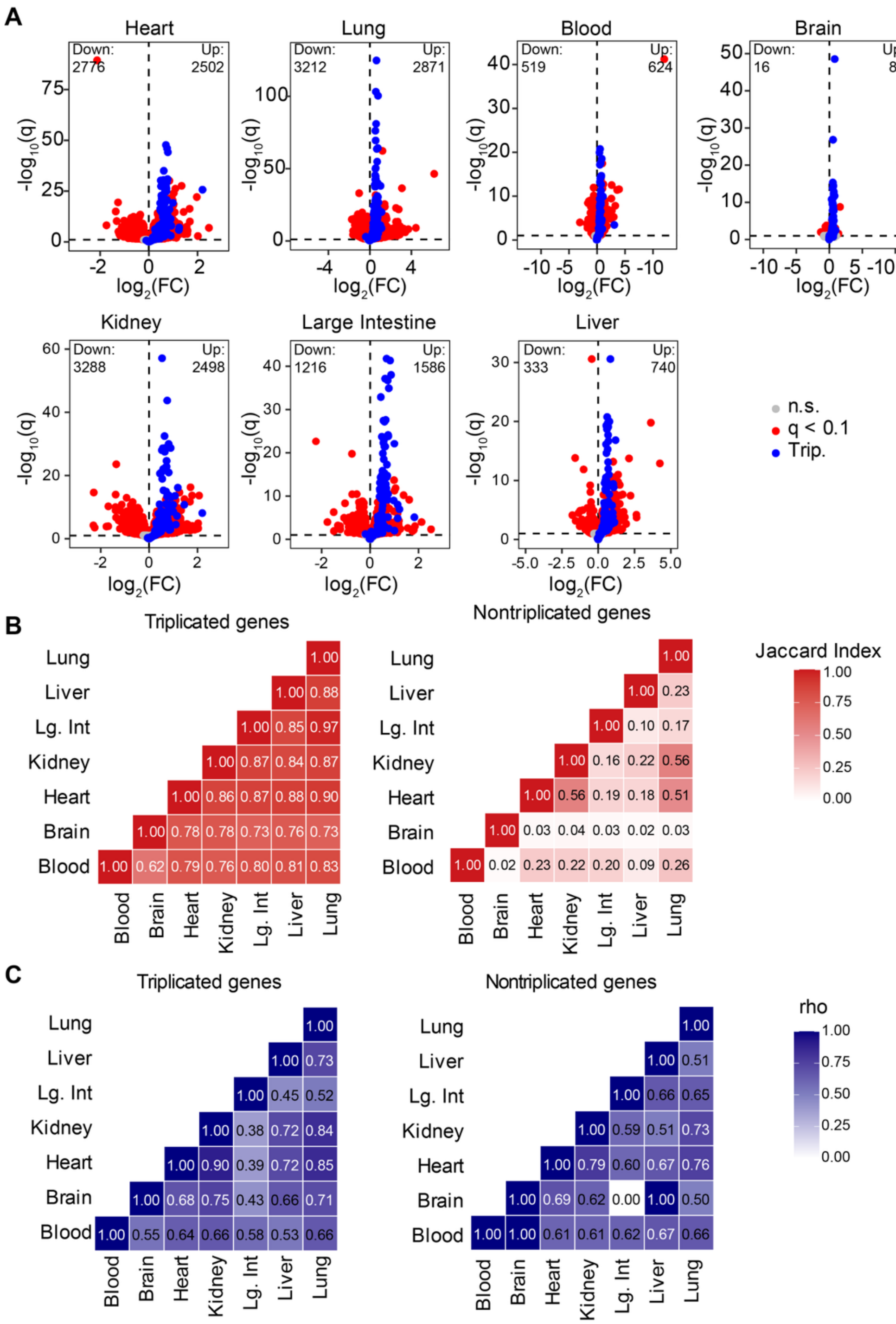


**Supplementary Figure 2, related to Figure 2: Tissue-specific differential expression at the 4-month timepoint. (A)** Volcano plots of differentially expressed genes in Dp16 vs. WT mice for each of seven tissues at 4 months (12 WT and 12 Dp16 mice per tissue). Triplicated genes are shown in blue, nontriplicated genes meeting the significance threshold (q<0.1) in red, and genes not meeting the threshold in grey. Note that axis scales differ between tissues. **(B)** Pairwise Jaccard similarity of differentially expressed genes between tissues, shown separately for triplicated (left) and nontriplicated (right) genes. **(C)** Pairwise Spearman rank correlations (rho) of per-gene $\log_2$ fold-changes between tissues, shown separately for triplicated (left) and nontriplicated (right) genes.

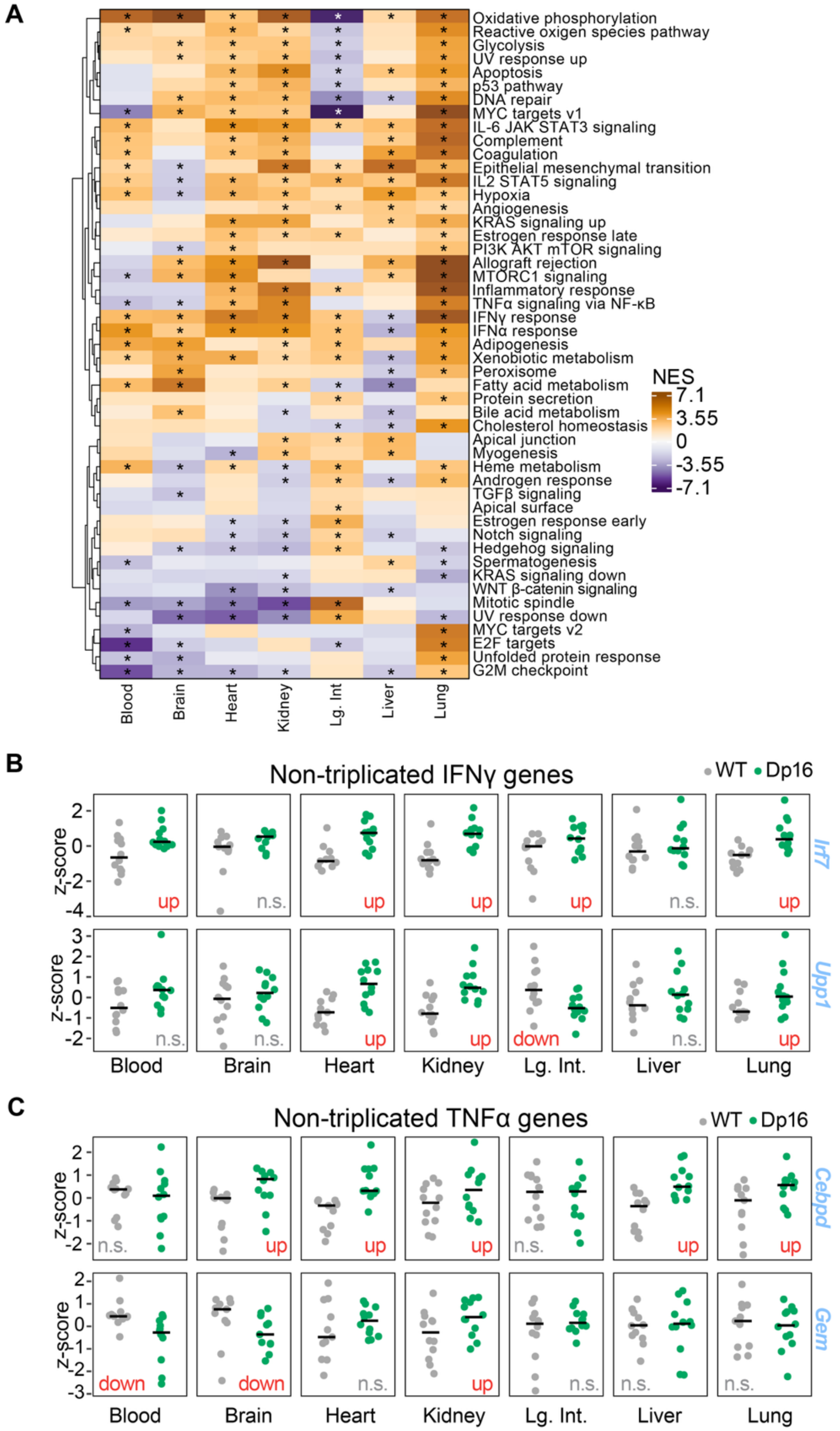


**Supplementary Figure 3, related to Figure 3. Inflammatory pathways are broadly enriched but composed of tissue-specific gene-level contributions at the 4-month timepoint. (A)** Heatmap of normalized enrichment scores (NES) from Gene Set Enrichment Analysis (GSEA) of Hallmark gene sets across 4-month tissues (12 WT and 12 Dp16 mice per tissue). Asterisks indicate q<0.1. **(B)** Expression of selected nontriplicated IFNγ response genes (*Irf7*, *Upp1*) in individual WT (grey) and Dp16 (green) mice across 4-month tissues. Values are z-score-transformed, SVA-adjusted $\log_2$ TPM; horizontal bars indicate the group mean. Genes are labeled up, down, or n.s. according to differential expression ($q<0.1$). **(C)** As in (B), for selected nontriplicated TNFα signaling via NFκB genes (*Cebpd*, *Gem*).

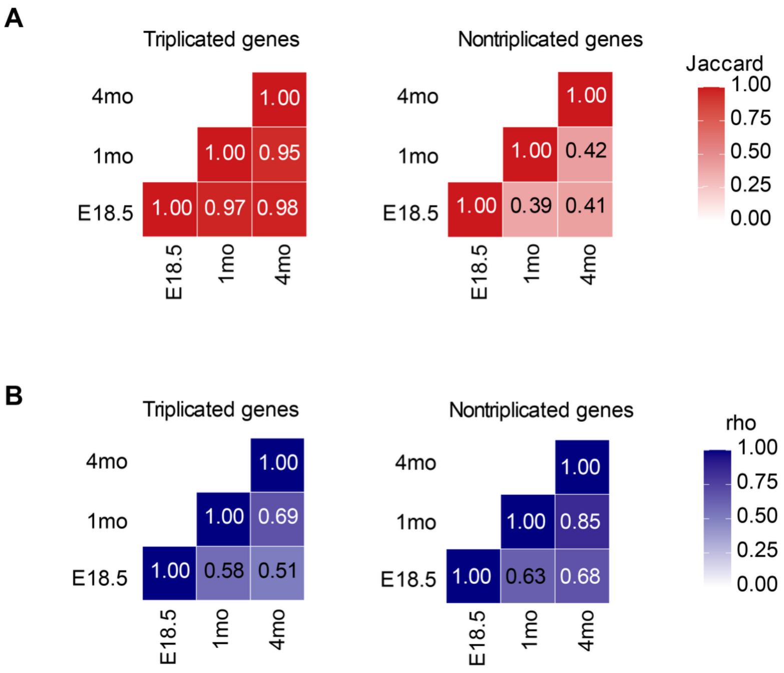


**Supplementary Figure 4, related to Figure 4. Developmental context shapes downstream transcriptional responses to Trisomy 21 in the lung. (A)** Pairwise Jaccard similarity of differentially expressed genes (DEGs) between lung timepoints, evaluated separately for triplicated (left) and nontriplicated (right) genes. 12 WT and 12 Dp16 mice per timepoint were evaluated. **(B)** Pairwise Spearman rank correlations (rho) for DEGs of per-gene $\log_2$ fold-changes between lung timepoints, evaluated separately for triplicated (left) and nontriplicated (right) genes.

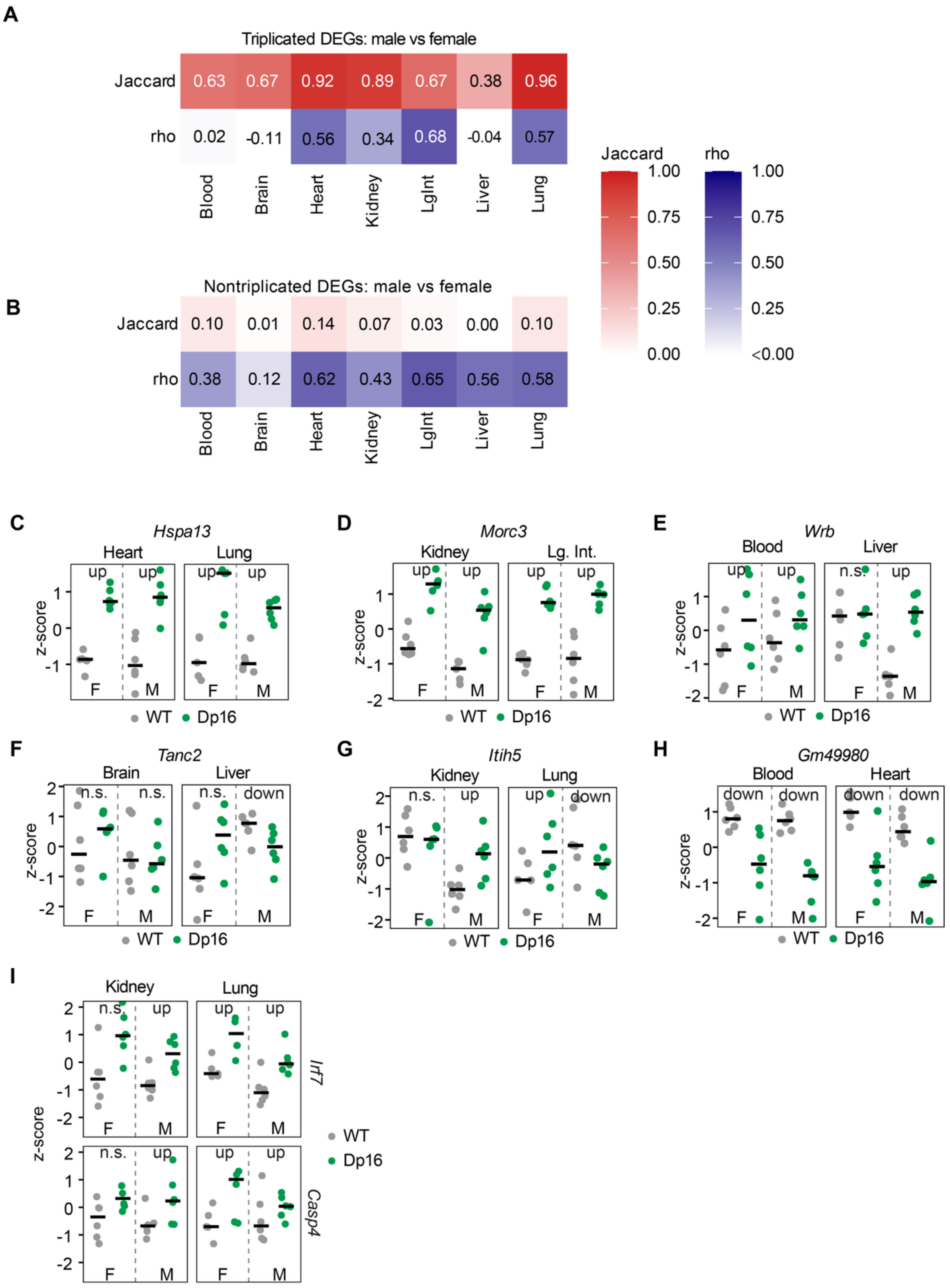


**Supplementary Figure 5, related to Figure 5. Sex modifies tissue-specific transcriptional responses to Trisomy 21 at 4 months. (A)** Pairwise Jaccard similarity (top) and Spearman rank correlation (rho, bottom) between male and female differentially expressed triplicated genes, by tissue. Six WT and six Dp16 mice were used per sex-timepoint-tissue group. **(B)** As in (A), for nontriplicated genes. **(C-I)** Sina plots of z-score-transformed, SVA-adjusted TPM expression for the genes shown in Fig. 5B, D, and F, displaying individual WT and Dp16 female (F) and male (M) mice. Genes are labeled up, down, or n.s. (not significant; q<0.1).

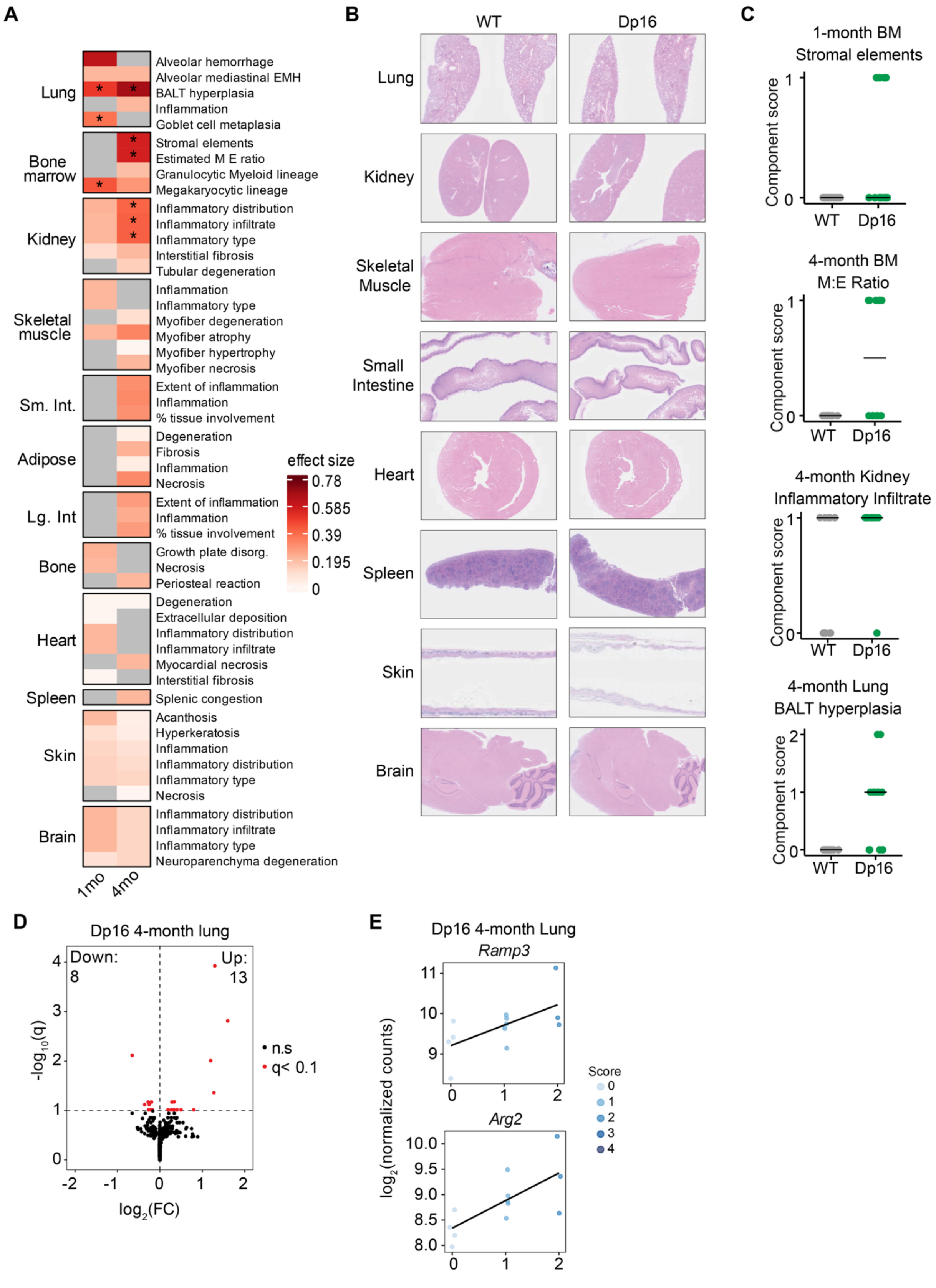


**Supplementary Figure 6, related to Figure 6. Integration of histopathology and transcriptomics links gene expression to tissue pathology. (A)** Heatmap of individual pathology component scores (effect size, Dp16 vs. WT) at 1 and 4 months across all tissues examined (12 WT and 12 Dp16 mice per tissue-timepoint). Grey indicates no pathology detected in either genotype or tissue not collected. **(B)** Representative histopathological images of WT and Dp16 tissues not shown in Fig. 6C. **(C)** Component scores for the four features highlighted in Fig. 6C. **(D)** Volcano plot of genes significantly associated ($q<0.1$) with composite pathology score in 4-month Dp16 lung, modeled using DESeq2 with composite score as a continuous covariate. **(E)** Expression of example hypoxia-related genes (*Ramp3*, *Arg2*) as a function of composite pathology score in 4-month Dp16 lung; point color indicates individual component score.

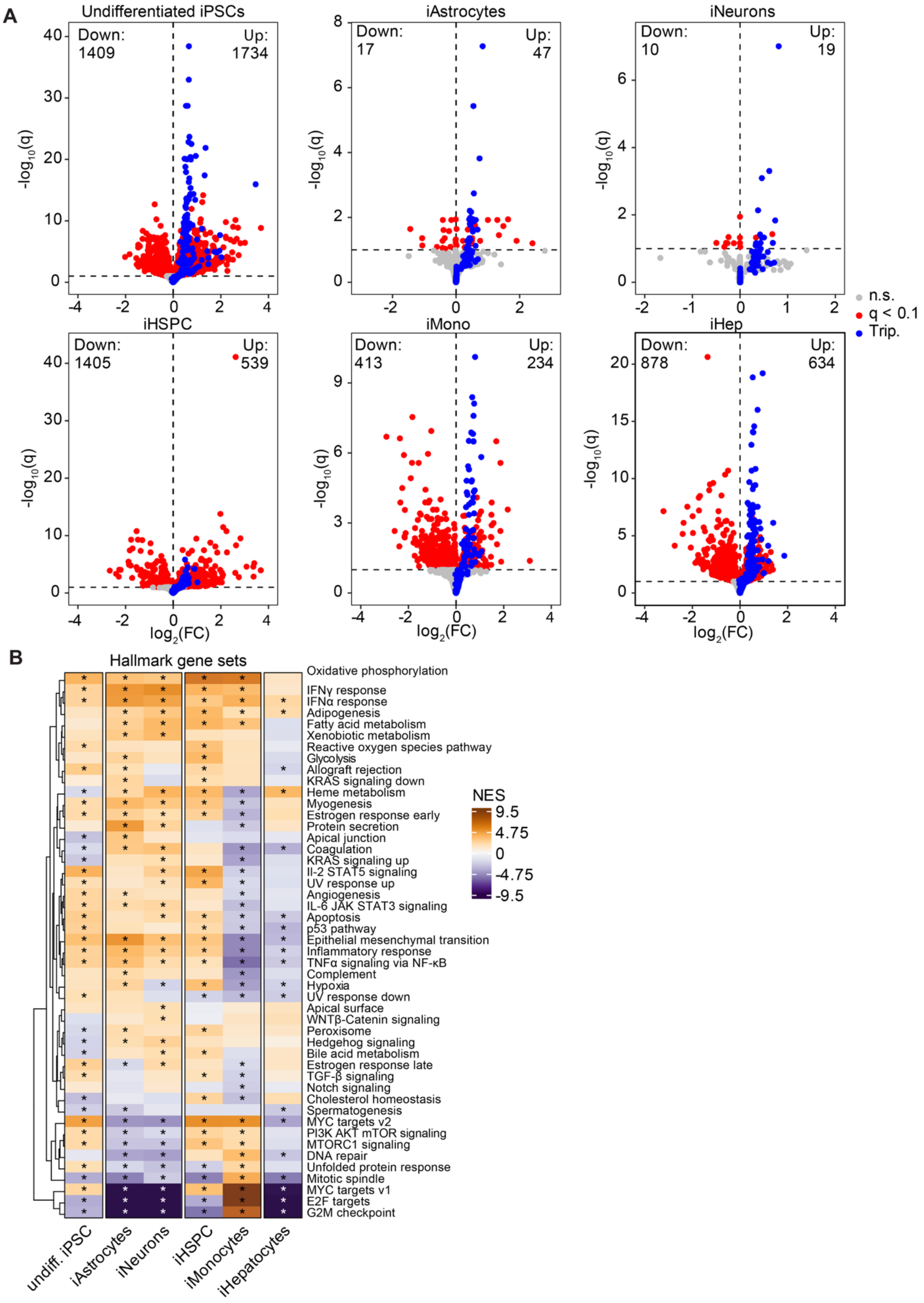


**Supplementary Figure 7, related to Figure 7. Human iPSC-derived cell types recapitulate conserved dosage effects and cell-type-specific transcriptional responses to Trisomy 21. (A)** Volcano plots of differentially expressed genes (T21 vs. D21, q<0.1) in each iPSC-derived cell type. Triplicated genes are shown in blue, nontriplicated genes meeting the significance threshold in red, and genes not meeting the threshold in grey. Cell types comprise 6 D21 and 6 T21 cell lines. **(B)** Heatmap of normalized enrichment scores (NES) from Gene Set Enrichment Analysis (GSEA) of all gene sets tested across cell types. Asterisks indicate $q<0.1$.
